# A lineage-specific *STAT5B*^N642H^ mouse model to study NK-cell leukemia

**DOI:** 10.1101/2023.10.04.560502

**Authors:** Klara Klein, Sebastian Kollmann, Julia List, Angela Hiesinger, Jonatan Kendler, Mehak Rhandawa, Jana Trifinopoulos, Barbara Maurer, Reinhard Grausenburger, Richard Moriggl, Thomas Rülicke, Agnieszka Witalisz-Siepracka, Wencke Walter, Gregor Hoermann, Veronika Sexl, Dagmar Gotthardt

## Abstract

Patients with T- and NK-cell neoplasms frequently have somatic *STAT5B* gain-of-function mutations. The most frequent *STAT5B* mutation is *STAT5B*^N642H^, which is known to drive murine T-cell leukemia although its role in NK-cell malignancies is unclear.

Introduction of the *STAT5B*^N642H^ mutation into human NK-cell lines enhances their potential to induce leukemia in mice. We have generated a mouse model that enables tissue-specific expression of *STAT5B*^N642H^ and have selectively expressed the mutated *STAT5B* in hematopoietic cells (N642H^vav/+^) or exclusively in NK cells (N642H^NK/NK^). All N642H^vav/+^ mice rapidly develop an aggressive T-/NK T-cell leukemia, whereas N642H^NK/NK^ mice display an indolent chronic lymphoproliferative disorder of NK cells (CLPD-NK) that progresses to an aggressive leukemia with age. Samples from NK-cell leukemia patients have a distinctive transcriptional signature driven by mutant STAT5B, which overlaps with that of murine *STAT5B*^N642H^-expressing NK cells.

We have generated the first reliable *STAT5B*^N642H^-driven pre-clinical mouse model that displays an indolent CLPD-NK progressing to aggressive NK-cell leukemia. This novel *in vivo* tool will enable us to explore the transition from an indolent to an aggressive disease and will thus permit the study of prevention and treatment options for NK-cell malignancies.

**Key points:**

- Generation of a lineage-specific *STAT5B*^N642H^ transgenic mouse model which develops NK-cell leukemia
- Leukemic NK cells with a STAT5B gain of function mutation have a unique transcriptional profile in mice and human patients

## Introduction

Natural Killer (NK)-cell malignancies are rare types of cancer that originate from the abnormal growth and proliferation of NK cells. They can be aggressive and challenging to treat. The World Health Organization distinguishes the following types of NK-cell neoplasms: extranodal NK/ T-cell lymphoma (ENKL), aggressive NK-cell leukemia (ANKL), chronic active Epstein-Barr virus (EBV) infection of NK cells (CAEBV) and CLPD-NK^1^. ENKL and ANKL are EBV-positive and associated with a poor prognosis^1^. CLPD-NK is usually EBV-negative, represents a subset of large granular lymphocytic leukemia (LGLL) and is largely an indolent disease that may develop into an aggressive NK-cell malignancy^1–6^. The factors that trigger the transformation of an indolent into an aggressive form of NK-cell leukemia are unknown.

Signal transducer and activator of transcription 5 (STAT5) is part of the Janus kinase (JAK)/STAT pathway and is crucial for the survival, proliferation and functionality of various hematopoietic cell types^7^. In leukemia, STAT activity is often enhanced by aberrant upstream tyrosine kinase activation, copy number gains^8^ or by activating mutations within the STAT3/5B proteins themselves^9–14^. STAT5 is the most frequently deregulated member of the JAK/STAT family in hematopoietic cancer^15–17^. It comprises two individual genes, *STAT5A* and *STAT5B*, which express proteins with high homology^7,18–20^. Although STAT5A and STAT5B have largely redundant functions, they both have some individual roles^18,21^. STAT5B is the dominant gene product in T and NK cells, in which it promotes their survival, proliferation, and cytotoxicity. *Stat5b*-deficient mice show reduced numbers of NK cells and humans with a STAT5B deficiency suffer from immunodeficiencies caused by impairment in T-, regulatory T-(T_reg_), and NK-cell differentiation and activation^18,22–31^.

Activating *STAT5* mutations in hematological cancer are almost exclusively found in *STAT5B*^12^. The most frequent *STAT5B* gain-of-function (GOF) mutation, *STAT5B*^N642H^, has been described in various forms of lymphoproliferative disorders, including T-cell lymphoma/leukemia, γδ-T-cell lymphoma, LGLL and NK-cell malignancies^9,11,13,14,32–42^. Activating *STAT5B* mutations have been detected in various NK-cell malignancies including cases of CLPD-NK and are linked to an aggressive clinical course^5,6,11,14,33,34,36,43^. *STAT5B*^N642H^ gives a proliferative advantage to human NK cells^36^, but whether it alone is sufficient to drive NK-cell leukemia is not known.

Compared to the wild type allele, *STAT5B*^N642H^ enhances dimer stability and causes elevated and prolonged STAT5B tyrosine phosphorylation. As a consequence, STAT5B target gene expression is increased^9–11,32,35,38^. But even in the presence of activating *STAT5B* mutations, upstream cytokine signaling is necessary to activate JAK/STAT5 signaling^10,36,44^. In immune cells, STAT5 signaling is induced by various cytokines, including interleukin (IL)-2, IL-7 and IL-15, which promote cell proliferation, survival and maturation^45,46^. IL-15 overexpression has been implicated in leukemogenesis as IL-15 transgenic mice develop NK- or NKT-cell leukemia^47–49^. Another transgenic mouse model expressing human IL-15 and a transgenic mouse model expressing human *STAT5B*^N642H^ in the hematopoietic system under the *vav* promoter (vav-N642H) develop a lethal CD8^+^ T-cell expansion^10,50^. The rapid development of the highly aggressive CD8^+^ T-cell disease may mask and prevent the development of other malignancies as transplantation of NKT or γδ T cells from vav-N642H mice lead to leukemia development^9,51^. To investigate the oncogenic potential of *STAT5B*^N642H^ in other cell types, we generated a lox-stop-lox *STAT5B*^N642H^ transgenic mouse model that enables lineage-restricted transgene expression driven by cell type-specific expression of the *Cre* recombinase. The use of *Cre* recombinase under an NK-cell specific promoter allows us to study the role of *STAT5B*^N642H^ in NK cells. Here, the transcriptional changes associated with *STAT5B*^N642H^ expression closely resemble the human STAT5B GOF disease signatures enabling the assessment of further targeted treatment options.

## Materials and Methods

### Conditional N642H mouse generation

Rosa26 (R26)-targeted lox-stop-lox (LSL) STAT5B and STAT5B^N642H^ knock-in mice were generated using a STOP-EGFP-ROSA-CAG (SERCA) targeting vector (obtained from Prof. Wunderlich, Uni Köln) integrating into the R26 locus and allowing transgene expression under the control of the CAG promoter coupled to internal ribosomal entry site (IRES)-controlled GFP expression upon Cre recombinase-mediated excision of the floxed STOP-cassette. C-terminally V5-tagged human *STAT5B* or *STAT5B*^N642H^ transgenes were cloned downstream of the STOP cassette into the SERCA targeting vector. A construct lacking a transgene and only having IRES-controlled EGFP downstream of the STOP cassette was included as a control. The three R26-LSL knock-in lines B6-*Gt(ROSA)26Sor*^*tm1(STAT5B-N642H)*^Biat, B6-*Gt(ROSA)26Sor*^*tm2(STAT5B)*^Biat and B6-*Gt(ROSA)26Sor*^*tm3(EGFP)*^Biat were generated using the linearized SERCA targeting vectors for electroporation into C57BL/6N embryonic stem (ES) cells (parental ES cell line C2, Stock Number: 011989-MU, Citation ID: RRID: MMRRC_011989-MU). Positively screened ES cell clones were injected into BALB/c blastocysts and transferred to pseudopregnant recipient mice. Chimeric offspring were bred with C57BL/6N mice and germ-line transmission was confirmed by PCR (forward primers: *5’-GCACTTGCTCTCCCAAAGTCGCTC-3’* (R26_wt_fw) and *5’-CGCCGACCACTACCAGCAGAACAC-3’* (R26_EGFP_fw); reverse primer: *5’-ACAACGCCCACACACCAGGTTAGC-3’* (R26_wt_rev)) and selected for further crossing to Cre lines.

### RNA-Seq

After 7 days of IL-2 culture, transgene-expressing NK cells (GFP^+^CD3^-^NK1.1^+^ cells isolated and cultured from GFP^NK/NK^, STAT5B^NK/NK^ or N642H^NK/NK^ mice) or control (Cre neg) NK cells (GFP^-^CD3^-^NK1.1^+^ cells isolated and cultured from Cre-negative N642H^STOP/STOP^ mice) were sorted with a BD FACS Aria III Sorter. Sorted NK cells were lysed in RLT buffer and RNA isolated using the RNeasy Mini kit (Qiagen). RNA-sequencing was performed at the Vienna BioCenter Core Facilities (VBCF). RNA-Seq libraries were prepared using the NEBNext Multiplex Oligos for Illumina (Dual Index Primers). Single-end, 50 bp sequencing was performed on an Illumina HiSeqV4 SR50 sequencer (Illumina, San Diego, CA, USA). After quality control of raw data with FastQC and removal of adapters and low-quality reads with Trimmomatic (version 0.36), reads were mapped to the GENECODE M13 genome using STAR (version 2.5.2b) with default parameters. Counts for union gene models were obtained using featureCounts from the Subread package (version 1.5.1). Differentially expressed (Benjamin–Hochbert corrected *p*-value (*p*-adjust) < 0.05 and fold change >2) genes were identified using edgeR (version 3.30.3). Over-representation analysis was done using the clusterProfiler (version 4.6.0). The RNA-Seq data reported in this article have been deposited in the Gene Expression Omnibus database (Accession ID: GSE236362).

### Human patient data

Primary samples were obtained from bone marrow (BM) or peripheral blood (PB) obtained at diagnosis or treatment from NK-cell neoplasms patients (n=64, from which three harbor activating *STAT5B* mutations (*STAT5B*^*N642H*^, *STAT5B*^*Q706L*^, *STAT5B*^*Y665F*^ and *STAT5B*^*V712E*^ mutations)) or healthy donors after informed consent. DNA and RNA were isolated from total leukocytes and whole-genome sequencing and RNA-sequencing were performed as described^52^ at the MLL Munich Leukemia Laboratory. Reads were aligned to the human reference genome (GRCh37, Ensembl annotation) using the Isaac aligner (v3.16.02.19) with default parameters. Tumor-unmatched normal variant calling was performed with a pool of sex-matched DNA (Promega, Madison, WI) using Strelka (v.2.4.7). Variants were queried against the gnomAD database (v.2.1.1) to remove common germline calls (global population frequency >1%) and annotated with Ensembl VEP. Analysis was restricted to protein-altering and canonical splice-site variants. For transcriptome analysis, the TruSeq Total Stranded RNA kit was used, starting with 250ng of total RNA, to generate RNA libraries following the manufacturer’s recommendations (Illumina, San Diego, CA, USA). 2x100bp paired-end reads were sequenced on the NovaSeq 6000 (Illumina, San Diego, CA, USA) with a median of 50 million reads per sample. Reads were mapped with STAR aligner (v2.5.0a) to the human reference genome hg19 (RefSeq annotation). Gene- and transcript-specific read abundance was calculated with Cufflinks (v2.2.1). For gene expression analysis, estimated read counts for each gene were normalized by Trimmed mean of M-values (TMM) normalization and the resulting log2 counts per million (CPMs) were used.

### Statistical analysis

The appropriate statistical method was used based on testing for normal distribution and homogeneity of variance. Tests were performed using GraphPad Prism. The statistical test is indicated in the corresponding figure legend.

### Data Sharing Statement

All other relevant data that support the conclusions of the study are available from the authors on request. Please contact

## Results

### N642H^vav/+^ mice develop a hematopoietic malignancy

We generated mice with a human V5-tagged *STAT5B*^N642H^ transgene and IRES-eGFP under the CAG promoter downstream of a lox-stop-lox-cassette integrated into the *Rosa26* locus. The animals were first crossed to *Vav*-Cre mice^32^ (N642H^vav/+^) to study the effects of *STAT5B*^N642H^ on the hematopoietic system (**Figure S1A**). We validated the presence of the STAT5B^N642H^ transgene by analyzing eGFP expression and the presence of the V5 tag in N642H^vav/+^ mice (**Figure 1A, S1B**). N642H^vav/+^ BM cells displayed increased levels of tyrosine-phosphorylated STAT5 (pYSTAT5) compared to control BM cells, although reduced levels when compared to the pYSTAT5 levels of vav-N642H mice^43^ (**Figure 1A**). N642H^vav/+^ mice at 8 weeks of age had an elevated BM cellularity and an enlarged hematopoietic stem cell (HSC) pool under homeostatic conditions (**Figure S1C-F**). Numbers of erythroid cells (Ter119^+^) and NK cells (CD3^-^NK1.1^+^) were reduced, while T cells (CD3^+^CD4^+^ or CD3^+^CD8^+^), B cells (CD19^+^) and myeloid cells (CD11b^+^Gr1^+^) were increased in the BM (**Figure S1G**). At 8 weeks of age, N642H^vav/+^ mice showed splenomegaly (**Figure 1B**) with significantly expanded myeloid and B-cell compartments (**Figure 1C, Figure S1H**). The peripheral blood of N642H^vav/+^ mice lacked any significant alterations in the composition of leukocytes, except for a decrease in the frequency of CD4^+^ T cells (**Figure S1I**).

**Figure 1.**
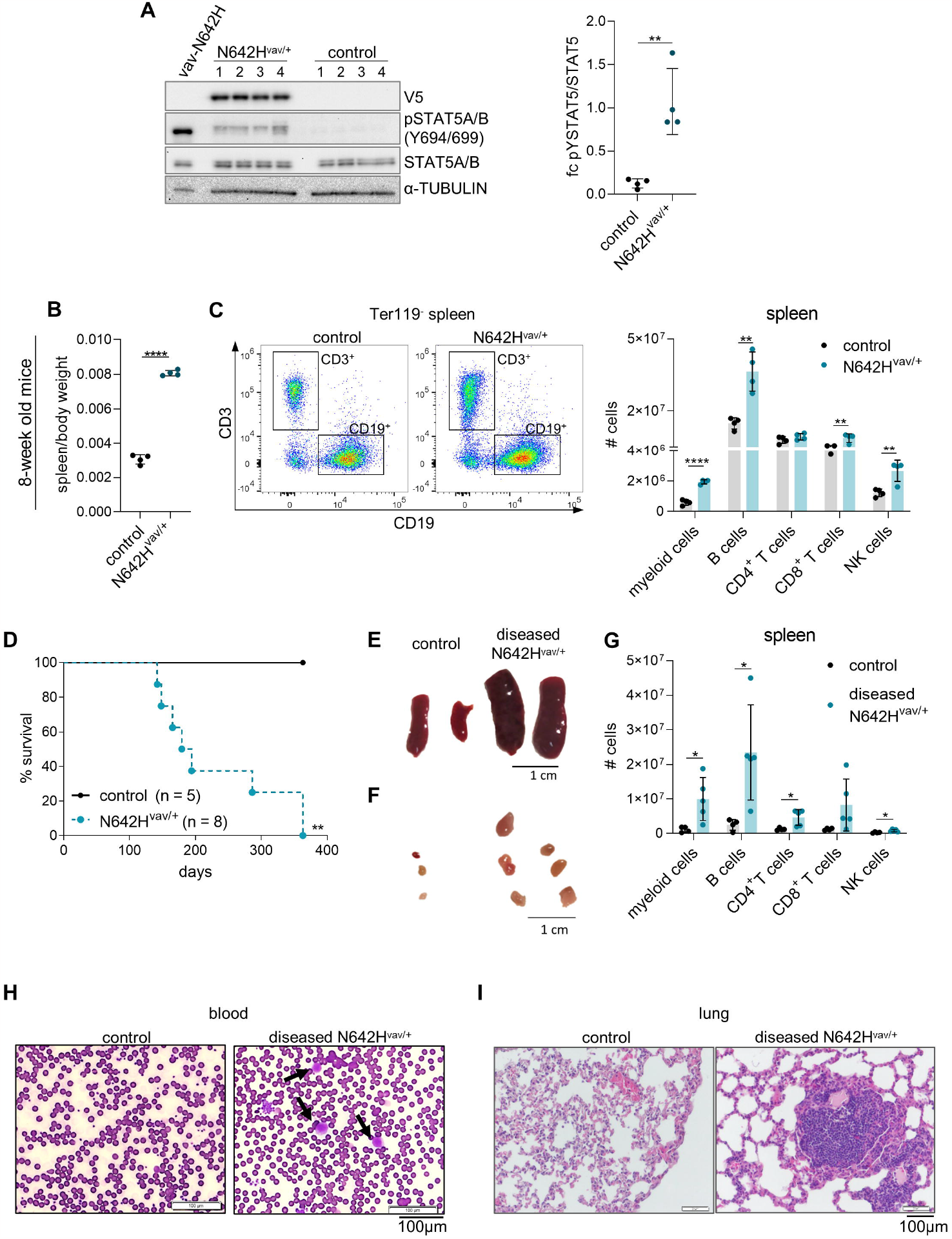
N642H^vav/+^ mice develop a hematopoietic malignancy. (A) (*Left*) V5, pYSTAT5, STAT5A/B and immunoblot analysis of BM from vav-N642H (n=1), control (n=4) and N642H^vav/+^ mice (n=4). α-TUBULIN served as loading control. (*Right*) Quantification of fold change (fc) of pYSTAT5 relative to STAT5A/B immunoblot levels from 8-week-od control and N642H^vav/+^ BM (n=4/genotype, mean±SD). (B) Relative quantification of spleen weights from 8-week-old control and N642H^vav/+^ mice (n=4/genotype, mean±SD). (C) (*Left*) Representative flow cytometric gating of CD3^+^ and CD19^+^ populations on Ter119 negative spleen cells and (*Right*) quantification of total splenic CD11b^+^Gr1^+^ myeloid cells, CD19^+^ B-cells, CD3^+^CD4^+^-, CD3^+^CD8^+^ T cells and CD3^-^NK1.1^+^ NK cells from 8-week-old control and N642H^vav/+^ mice (n=4/genotype, mean±SD). (D) Survival analysis of N642H^vav/+^ and control mice (n=5-8/genotype). Representative pictures of (E) SPs and (F) LNs of aged control and diseased N642H^vav/+^ mice. (G) Quantification of total splenic CD11b^+^Gr1^+^ myeloid cells, CD19^+^ B cells, CD3^+^CD4^+^-, CD3^+^CD8^+^ T cells and CD3^-^NK1.1^+^ NK cells from control and diseased N642H^vav/+^ mice (n=5/genotype, mean±SD). (H) Hemacolor® Rapid staining of blood smears from control and diseased N642H^vav/+^ mice (one representative picture/genotype, n=4/genotype). (I) H&E staining on lung tissue from control and diseased N642H^vav/+^ mice (one representative picture/genotype, n=4/genotype). Levels of significance were calculated using unpaired t-test in (A) - (C) and (G) and Mantel-Cox text in (D). *p < 0.05, **p < 0.01 and ****p < 0.0001.

Upon ageing, all N642H^vav/+^ mice developed a hematopoietic malignancy with a median survival of 186 days (**Figure 1D**). The mice suffered from reduced body weight and enlarged spleen and lymph nodes (**Figure 1E+F, S1J+K**). They had significantly elevated numbers of mature hematopoietic cell types in spleen, blood and lymph nodes but not in the BM (**Figure 1G, S1L-N**). Cell numbers were elevated in all lineages and no particular cell type was dominantly expanded (**Figure S1O**). Blood smears of the diseased N642H^vav/+^ mice showed leukemic blast-like cells (**Figure 1H**). The immune cell infiltration in the lungs was associated with a disruption of the regular lung architecture (**Figure 1I**).

### N642H^vav/+^ splenocytes drive the expansion of leukemic T-/NKT cells upon transplantation

To test whether the hematopoietic malignancy is transplantable, we injected Ly5.2^+^ splenic cells of diseased N642H^vav/+^ and healthy control mice into immunodeficient NSG recipients (NOD.Cg-Prkdc^scid^ Il2rg^tm1Wj1^/SzJ). All recipients of N642H^vav/+^ splenocytes developed a disease within 3 months (**Figure 2A**). Ly5.2^+^ N642H^vav/+^ cells densely infiltrated the BM, spleen and lung of the NSG recipients, indicating development of a leukemia (**Figure 2B, S2A-C)**. The infiltrating cell types were either T or NKT cells (**Figure 2C+D**). N642H^vav/+^ T and NKT cells expressed almost exclusively TCRβ but not TCRγδ (**Figure 2E+F)**. CD4^+^ and CD8^+^ T/NKT cells expanded in the recipient mice (**Figure 2E+G**). This argues against the idea that a specific T/NKT cell subtype is driving the leukemia. The diseases that develop in N642H^vav/+^ mice closely resemble the T-/NKT-cell diseases observed in patients harboring the *STAT5B*^*N642H*^ mutation^9^.

**Figure 2.**
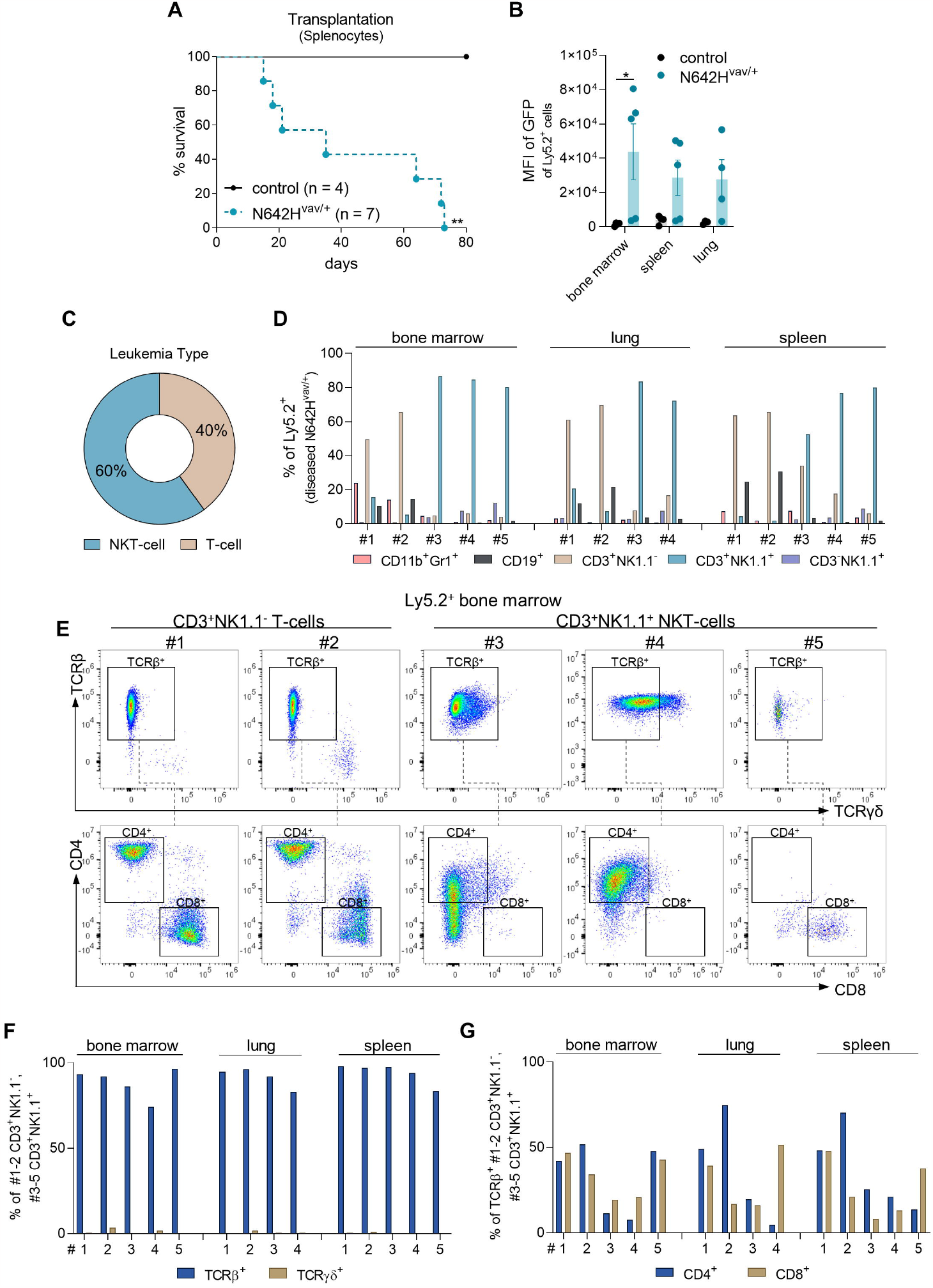
*STAT5B*^*N642H*^ drives the expansion of leukemic NKT-/T cells upon transplantation. Splenic cells (Ly5.2^+^) from either control or diseased N642H^vav/+^ mice were intravenously (*i*.*v*.) injected into NSG mice and analyzed. (A) Survival analysis (n=4-7/genotype). (B) Quantification of GFP levels among transplanted Ly5.2^+^ cells in BM, spleen and lung of NSG mice injected with control or N642H^vav/+^ cells (n=3-5/genotype, mean±SD). (C) Quantification of the leukemia type developed by the N642H^vav/+^ transplanted NSG recipients. (D) Relative quantification of injected Ly5.2^+^ N642H^vav/+^ CD11b^+^Gr1^+^ myeloid cells, CD19^+^ B cells, CD3^+^NK1.1^-^ T cells, CD3^+^NK1.1^+^ NKT cells and CD3^-^NK1.1^+^ NK cells in BM, lung and spleen of NSG mice (n=4-5). (E) Representative FACS plots of TCRβ and TCRγδ gating starting from CD3^+^NK1.1^-^ T-cells or CD3^+^NK1.1^+^ NKT-cells in BM of N642H^vav/+^ transplanted NSG mice. (F) Relative quantification of TCRβ or TCRγδ expression on injected Ly5.2^+^CD3^+^NK1.1^-^ or Ly5.2^+^CD3^+^NK1.1^+^ N642H^vav/+^ cells (n=4-5). (G) Relative quantification of CD4 or CD8 expression on Ly5.2^+^TCRβ^+^CD3^+^NK1.1^-^ or Ly5.2^+^ TCRβ^+^CD3^+^NK1.1^+^ N642H^vav/+^ injected cells (n=4-5). Levels of significance were calculated using Mantel-Cox text in (A) and Mann-Whitney test in (B). *p < 0.05 and **p < 0.01.

### *STAT5B*^N642H^ promotes cytokine independence of human NK-cell lines

Despite the presence of an activating STAT5B mutation, N642H^vav/+^ mice did not develop NK-cell leukemia. To investigate the oncogenic potential of STAT5B^N642H^ in NK cells, we ectopically expressed human STAT5B or STATB^N642H^ in two human NK-cell lines (IMC-1 and KHYG-1) that harbor *TP53* mutations but lack mutations in the JAK/STAT3/5 pathway^33,53^. Transduction with STAT5B or STAT5B^N642H^ decreased cell growth in standard IL-2 culture (100U/ml) (**Figure S3A+B**) but gave a growth advantage at limited IL-2 concentrations (25U/ml) (**Figure S3C+D**). In the absence of IL-2, STAT5B^N642H^ was required for cytokine-independent growth (**Figure 3A+B, Figure S3E+F**). This prompted us to test whether STAT5B^N642H^ enhances the disease-initiating potential of KHYG-1 and IMC-1 cells *in vivo* (**Figure 3C**). When injected into NSG mice, STAT5B^N642H^-expressing IMC-1 cells accelerated disease onset significantly compared to parental and non-mutant STAT5B-expressing cells (**Figure 3D-G**). In contrast, neither the parental nor the STAT5B-overexpressing KHYG-1 cells caused disease in NSG recipient mice. All STAT5B^N642H^-transduced KHYG1 cells induced leukemia within 21-25 days (**Figure 3D**). The disease primarily manifested in the BM and the liver (**Figure 3E-G**), both typical sites of disease manifestation in NK-cell leukemia patients^35,40,41,55^.

**Figure 3.**
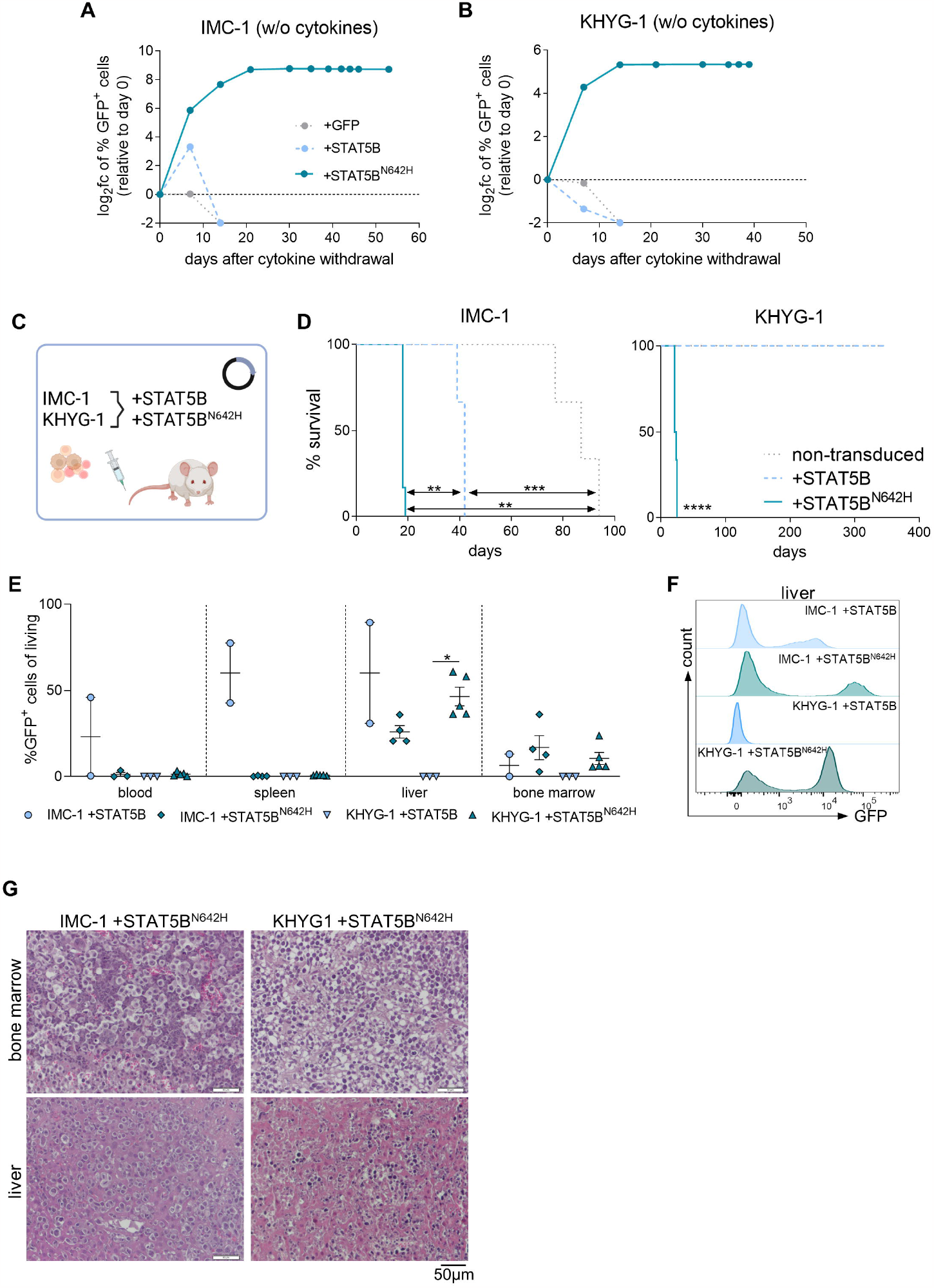
*STAT5B*^*N642H*^ promotes cytokine independence of leukemic human NK cells. (A) IMC-1 and (B) KHYG-1 cell lines were transduced with non-mutant STAT5B (+STAT5B) or STAT5B^N642H^ (+STAT5B^N642H^). As a control, cells were transduced with the empty vector, carrying only IRES-controlled eGFP (+GFP). After initial culture in presence of IL-2, transduced cells were completely deprived of IL-2. The percentage of transduced (GFP^+^) cells depicted as log_2_ fc relative to day 0 was monitored over time after cytokine withdrawal. (C) Schematic overview for transplantation of cytokine-independent STAT5B^N642H^ transduced, IL-2 dependent non-mutant STAT5B transduced and non-transduced IMC-1 and KHYG-1 cells into immunodeficient recipient mice. (D) Survival analysis of IMC-1 (*Left*) and KHYG-1 (*Right*) transplanted NSG mice (n = 3-6/cell line). (E) Flow cytometric analysis of percentage of transplanted non-mutant STAT5B or STAT5B^N642H^ transduced (GFP+) cells among all living cells in the indicated organs (n=2-5/genotype, mean±SD). (F) Representative histograms for GFP signal within living cells in the liver of transplanted mice. (G) Representative images of H&E-stained BM and liver tissue from NSG mice transplanted with STAT5B^N642H^ transduced (*Left*) IMC-1 or (*Right*) KHYG-1 cells. Levels of significance were calculated using Mantel-Cox text in (D) and Mann-Whitney test in (E). *p < 0.05, **p < 0.01, ***p < 0.001, ****p < 0.0001.

### An NKp46^+^-cell specific mouse model to study the oncogenic role of *STAT5B*^N642H^ in NK cells

To investigate the oncogenic role of *STAT5B*^N642H^ in NK cells in detail, we crossed the B6-*Gt(ROSA)26Sor*^*tm1(STAT5B-N642H)*^ mice to *Ncr1*-iCreTg mice^54^ (N642H^NK/NK^). These mice express *STAT5B*^N642H^ exclusively in NKp46^+^ cells, which mainly represent NK cells. A human *STAT5B* transgene-expressing mouse strain (STAT5B^NK/NK^) and a strain solely expressing eGFP (GFP^NK/NK^) were used as controls (**Figure 4A**). All *Cre*-positive litters expressed GFP in NK cells (**Figure S4A**) and we confirmed the V5-tagged transgene expression in STAT5B^NK/NK^ and N642H^NK/NK^ NK cells (**Figure 4B**).

**Figure 4.**
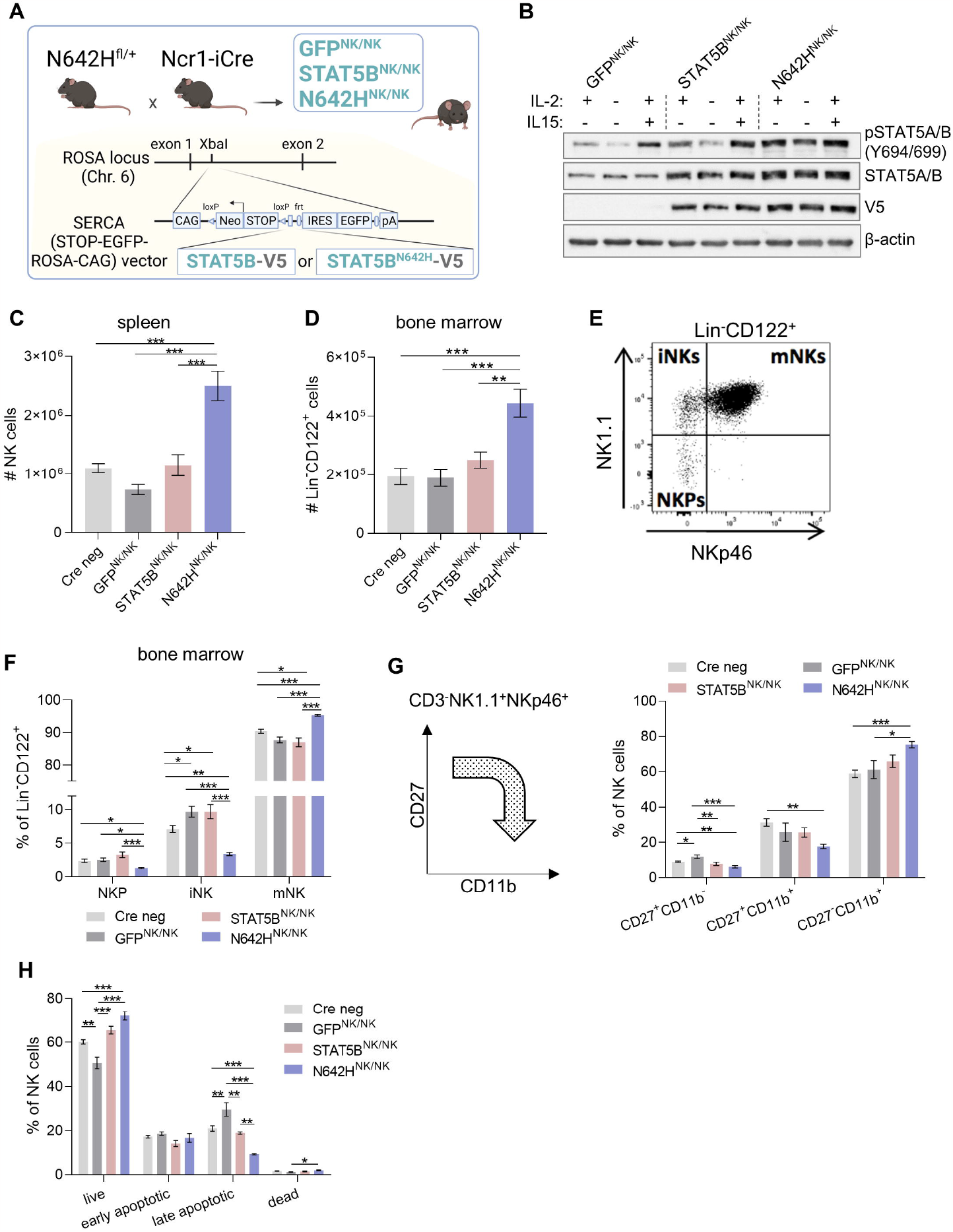
A NKp46^+^-cell specific mouse model to study the oncogenic role of *STAT5B*^N642H^ in NK cells. (A) Schematic overview of the generation of N642H^NK/NK^, STAT5B^NK/NK^ and GFP^NK/NK^ mice. (B) pYSTAT5, STAT5A/B and V5 immunoblot analysis of IL-2 cultured NK cells from GFP^NK/NK^, STAT5B^NK/NK^ and N642H^NK/NK^ mice. NK cells were either directly lysed, lysed after being starved off IL-2 for 3h or lysed after IL-2 starvation and restimulation with IL-2 and IL-15. β-Actin served as a loading control. (C) Absolute numbers of splenic NK cells (CD3^-^ NK1.1^+^NKp46^+^) in 8-12-week-old GFP^NK/NK^, STAT5B^NK/NK^, N642H^NK/NK^ and Cre negative (neg) control mice (n≥6/genotype, mean±SD). (D) Absolute numbers of NK cells (lineage (Lin) negative (CD3^-^CD19^-^Gr1^-^Ter119^-^) CD122^+^ cells) in BM of 8-12-week-old GFP^NK/NK^, STAT5B^NK/NK^, N642H^NK/NK^ and Cre neg mice (n=6-10/genotype, mean±SD). (E) Representative gating of NK-cell developmental stages among Lin-CD122^+^ cells within the BM, including NK1.1^-^NKp46^-^ NK cell precursors (NKPs), NK1.1^+^NKp46^-^ immature (iNKs) and NK1.1^+^NKp46^+^ mature NK cells (mNKs). (F) Percentages of NKPs, iNKs and mNKs among Lin^-^CD122^+^ BM cells (n≥10/genotype, mean±SD). (G) *(Left*) Schematic overview on splenic NK cell maturation stages based on the CD27 and CD11b expression. *(Right)* Percentages of CD27^+^CD11b^-^, CD27^+^CD11b^+^ and CD27^-^CD11b^+^ NK cells in the spleens of GFP^NK/NK^, STAT5B^NK/NK^, N642H^NK/NK^ and Cre neg mice (n≥6/genotype, mean±SD). (H) Apoptosis staining of splenic NK cells from GFP^NK/NK^, STAT5B^NK/NK^, N642H^NK/NK^ and Cre neg mice (n≥4/genotype, mean±SD). Levels of significance were calculated using one-way ANOVA in (C) – (H). *p < 0.05, **p < 0.01, ***p < 0.001, ****p < 0.0001.

STAT5B^N642H^ molecules have an enhanced capacity for self-dimerization and a reduced susceptibility to inactivation by dephosphorylation^9^. N642H^NK/NK^ splenic NK cells displayed enhanced pYSTAT5 levels upon IL-15 stimulation *ex vivo* (**Figure S4B+C**). STAT5B^NK/NK^ and N642H^NK/NK^ NK cells had elevated STAT5 protein levels compared to GFP^NK/NK^ NK cells. Upon IL-2 withdrawal, pYSTAT5 dephosphorylation was delayed in N642H^NK/NK^ NK cells (**Figure 4B)**.

N642H^NK/NK^ mice display increased NK-cell numbers (**Figure 4C+D, S4D+E**) and more mature NK cells in the BM and spleen compared to control strains (**Figure 4E-G, S4F**). Furthermore, *STAT5B*^N642H^ expression in NK cells was associated with increased survival and reduced apoptosis *ex vivo* (**Figure 4H**). The data are consistent with the idea that STAT5B promotes NK-cell survival and maturation^55^. We found enhanced levels of Granzyme B and Perforin in N642H^NK/NK^ NK cells (**Figure S4G**), supporting the role of STAT5B in regulating the levels of cytolytic molecules^22,24,31,55^.

### Conditional N642H^NK/NK^ mice develop NK-cell lineage leukemia

The oncogenic potential of *STAT5B*^N642H^ in NK cells was assessed by ageing of the animals. While the majority of the N642H^NK/NK^ mice had an indolent expansion of NK cells, around 30% (8/27) of them developed disease symptoms within 17 months, whereas only 2 of 50 control mice became moribund. One STAT5B^NK/NK^ and one Cre-negative control mouse were sacrificed after 486 and 518 days respectively due to unspecific age-related disease symptoms without any signs of leukemia (**Figure 5A**). The diseased N642H^NK/NK^ mice consistently displayed a leukemic phenotype and suffered from significant body weight loss, splenomegaly, and an expansion of GFP^+^ cells in various organs, including spleen, liver, BM and blood (**Figure 5B+C, Figure S5A+B**). NK cells (CD3^-^ TCR^-^NK1.1^+^) were the predominant cell type in the expanded GFP+ cells in the spleen of five out of eight diseased N642H^NK/NK^ mice. One of the diseased mice displayed an expansion of NK1.1^+^ γδ T cells, while two other mice had a predominant expansion of GFP^+^ lacking both NK- and T-cell markers (CD3^-^ TCR^-^NK1.1^-^ NKp46^-^ cells) (**Figure 5D+E, Figure S5C**). To test whether the leukemia-initiating cells belong to the NK-cell compartment, we transplanted splenic cells from the diseased N642H^NK/NK^ mice into NSG mice (**Figure 5F**). Transplantation initiated a fast-progressing disease. The development of NK-cell leukemia was confirmed in ∼70% of transplanted mice (**Figure 5G+H**), which showed hepatosplenomegaly, anemia (**data not shown**) and infiltration of BM, spleen, blood and liver (**Figure 5I, Figure S5D-K**). The transplantation of splenocytes from the mouse that had a lethal expansion of CD3^+^NK1.1^+^ TCRγδ^+^ T cells (#8) verified a disease driven by *STAT5B*^N642H^-expressing γδ T cells. The transplantation of splenic cells with an accumulation of GFP^+^ CD3^-^ TCR^-^NK1.1^-^ NKp46^-^ cells (#6 and #7) suggested that the mice suffered more likely from an NK-cell leukemia than an acute leukemia of T-cell origin as there was a pronounced NK1.1^+^ but not a CD3^+^ or TCR^+^ population in various organs upon transplantation (**Figure S5D-K**). We observed leukemic blast-like cells in the blood of all diseased recipient mice (**Figure 5I**) In summary, our data show that N642H^NK/NK^ mice predominantly develop a transplantable NK-cell leukemia.

**Figure 5.**
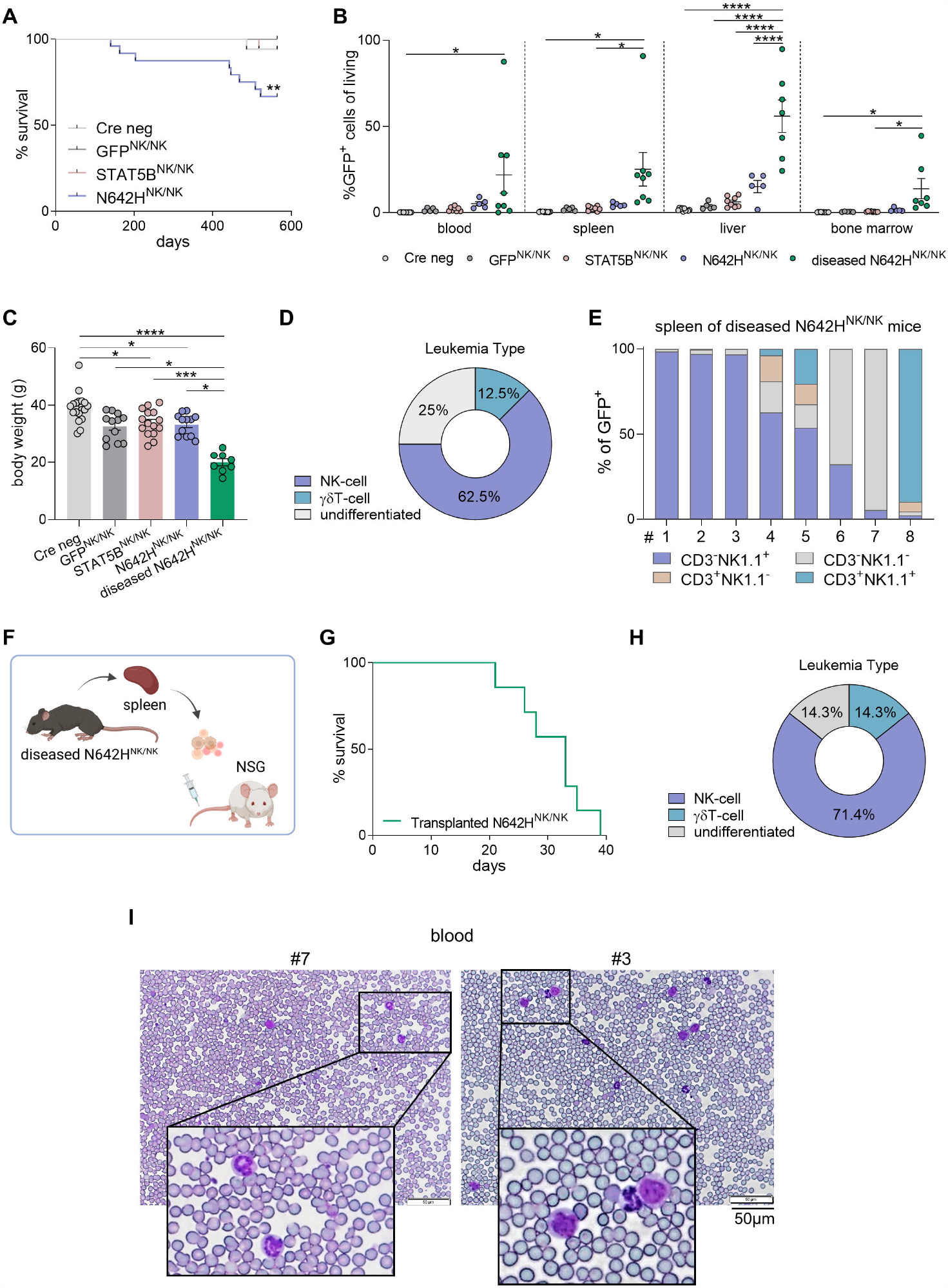
*STAT5B*^N642H^ induces NK-cell leukemia in mice. Cre neg, GFP^NK/NK^, STAT5B^NK/NK^ and N642H^NK/NK^ mice were aged and monitored for signs of disease development. (A) Survival analysis of Cre neg, GFP^NK/NK^, STAT5B^NK/NK^ and N642H^NK/NK^ mice (n≥10/genotype). (B) Flow cytometric analysis of GFP^+^ cells in different tissues of Cre neg, GFP^NK/NK^, STAT5B^NK/NK^, healthy N642H^NK/NK^ and diseased N642H^NK/NK^ mice (n≥5/genotype, mean±SD). (C) Body weight quantification of Cre neg, GFP^NK/NK^, STAT5B^NK/NK^, healthy N642H^NK/NK^ and diseased N642H^NK/NK^ mice (n≥8/genotype, mean±SD). (D) Quantification of the leukemia type developed by the N642H^NK/NK^ mice. (E) Relative quantification of CD3^+^NK1.1^-^ T cells, CD3^+^NK1.1^+^ NKT cells, CD3^-^NK1.1^+^ NK and CD3^-^NK1.1^-^ cells among GFP^+^ cells in spleen of diseased N642H^NK/NK^ mice (n=8). (F) Schematic overview of the *i*.*v*. transplantation of splenocytes from diseased N642H^NK/NK^ mice into NSG mice. (G) Survival analysis of transplanted NSG mice (n=7). (H) Quantification of the leukemia type developed by the NSG recipients of diseased N642H^NK/NK^ splenocytes. (I) Representative images of H&E-stained blood smears from NSG mice transplanted with diseased N642H^NK/NK^ splenocytes (*Left*) #7 and (*Right*) #3. Levels of significance were calculated using Mantel-Cox test in (A) and one-way ANOVA in (B) and (C). *p < 0.05, **p < 0.01, ***p < 0.001, ****p < 0.0001.

### N642H^NK/NK^ NK cells display molecular features of NK-cell leukemia patients with *STAT5B* GOF mutations

To investigate the molecular mechanism of *STAT5B*^N642H^-driven NK-cell leukemia, we performed RNA-Seq of IL-2-activated N642H^NK/NK^ and control (Cre neg, GFP^NK/NK^, STAT5B^NK/NK^) NK cells. All genotypes had different transcriptional profiles (**Figure S6A**). There was a higher number of significantly differentially expressed genes (DEGs) in N642H^NK/NK^ than in STAT5B^NK/NK^ NK cells (**Figure S6B+C**). Several biological pathways were commonly enriched in both STAT5B^NK/NK^ and N642H^NK/NK^ NK cells, albeit to varying degrees (**Figure S6D**). The *STAT5B*^N642H^ mutation enhances pYSTAT5B stability, promoting the expression of STAT5B target genes^9–11,32,35,38^. In N642H^NK/NK^ NK cells, we observed a predominant enrichment of IL-2-STAT5 and IL-6-STAT3 signaling pathways (**Figure S6D)**. To determine whether the transcriptional patterns induced by *STAT5B*^N642H^ in N642H^NK/NK^ NK cells resemble those found in NK-cell leukemia patients with *STAT5B* GOF mutations, we analyzed RNA-Seq data sets from 64 NK-cell leukemia patients. NK-cell leukemias show a different transcriptional profile than BCP-ALL, T-ALL and healthy controls (**Figure 6A**). Of the patients studied, three had *STAT5B* GOF mutations and 17 harbored mutations in other members of the JAK/STAT pathway. Comparison of the *STAT5B*^N642H^-specific DEGs (N642H^NK/NK^ vs. STAT5B^NK/NK^) from the murine NK cells with DEGs associated with *STAT5B* GOF mutations in human NK-cell leukemia patients (*STAT5B* GOF mutant vs. JAK/STAT non-mutant)^58,59^ identified a set of 239 common DEGs induced by mutant *STAT5B* (**Figure 6B; Table S1**).

**Figure 6.**
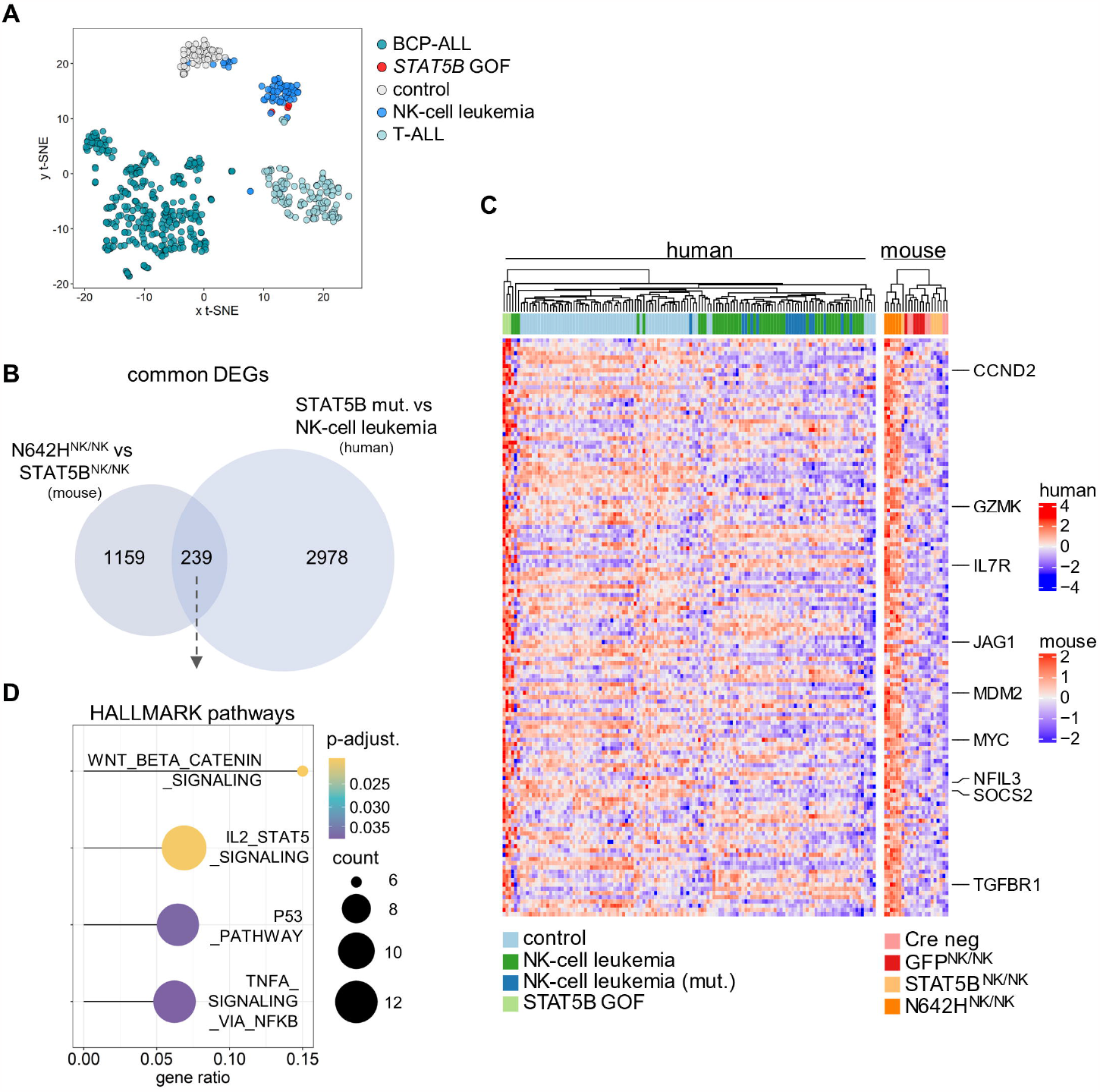
N642H^NK/NK^ NK cells display molecular features of NK-cell leukemia patients harboring *STAT5B* GOF mutations. RNA-sequencing data of NK cells from Cre neg, GFP^NK/NK^, STAT5B^NK/NK^, N642H^NK/NK^ mice and NK-cell leukemia patients was analysed. (A) PCA analysis of RNA-sequencing data of control, BCP-ALL, NK-cell leukemia, T-ALL and *STAT5B* GOF mutated NK-cell leukemia patient samples. (B) Venn diagram illustrating common DEGs from N642H^NK/NK^ vs. STAT5B^NK/NK^ and *STAT5B* GOF mut. vs. NK-cell leukemia samples. (C) Heatmap illustrating expression of the common upregulated genes (N642H^NK/NK^ vs. STAT5B^NK/NK^ and *STAT5B* GOF vs. NK-cell leukemia) of human control (n=64), NK-cell leukemia (n=44), NK-cell leukemia (mut., carrying a JAK/STAT mutation) (n=17), *STAT5B* GOF mut. (n=3) samples and murine Cre neg (n=6), GFP^NK/NK^ (n=5), STAT5B^NK/NK^ (n=5), N642H^NK/NK^ (n=6) NK cells. (D) Enriched HALLMARK pathways of the common DEGs found in (B).

Of these 239 *STAT5B* GOF DEGs, 136 were commonly upregulated and 103 commonly downregulated (**Figure 6C, Figure S6E**). The upregulated DEGs included known STAT5-regulated genes such as *SOCS2, CCND2, NFIL3* and *MYC*^24,55,56^ and several genes involved in signaling pathways and tumorigenesis (**Figure 6C**). We found WNT, IL2-STAT5, p53 and TNFα signaling significantly upregulated, but no pathways were significantly downregulated (p-adj ≤0.05) (**Figure 6D**).

In summary, our findings show that N642H^NK/NK^ NK cells exhibit transcriptional patterns resembling those found in *STAT5B*-mutated human NK-cell leukemia underlining the translational validity of the disease model. Hence, the genes identified to be regulated by *STAT5B*^N642H^ represent potential targets for novel therapeutic strategies in STAT5B-driven NK-cell malignancies.

## Discussion

STAT5B is a prominent driver of hematopoietic diseases^11^. The *STAT5B*^N642H^ mutation is primarily found in diseases arising from T/NKT cells^27^. Previously, a vav-STAT5B^N642H^ mouse model was reported to develop an aggressive CD8^+^ T-cell lymphoma^10^. We now describe a different mouse model (N642H^vav/+^ mice, **Figure 1-2**) that develops a slowly progressing CD4^+^, CD8^+^ T-cell or NKT-cell leukemia and succumbs to the disease within a year. N642H^vav/+^ mice display lower pYSTAT5 levels than vav-N642H mice. The different pYSTAT5 levels could stem from a difference in the levels of transgene expression or might reflect the more progressive CD8^+^ T-cell disease in young vav-N642H mice. The difference in disease type and onset might also be attributed to the different promoters and their regulatory environment that drives transgene expression (*CAG* vs. *Vav1*) in the individual cell types. N642H^vav/+^ mice not only develop a CD8^+^ T-cell leukemia but also display diverse disease phenotypes, making them a closer representation of patients with *STAT5B* GOF mutations^9^. It therefore represents an excellent model to study the impact of *STAT5B*^N642H^ in different cellular and disease contexts.

Our mouse model allows for lineage- or tissue-specific transgene expression enabling us to decipher STAT5B^N642H^’s functions in NK cells in more detail. *STAT5B*^N642H^ expression in the NK-cell lineage results in hyperactive STAT5B signaling in NK cells, elevated cell numbers, decreased apoptosis, increased maturation and higher levels of lytic granzymes. The persistently increased counts of mature NK cells in 8-12-week-old N642H^NK/NK^ mice, which do not display disease symptoms, is indicative of an indolent CLPD-NK phenotype^57,58^. This finding aligns with the indolent phenotype of CD4^+^ T-LGLL patients carrying *STAT5B* mutations^12^. One third of the N642H^NK/NK^ mice go on to develop an aggressive disease, suggesting that indolent cases of NK-cell malignancies can transform into aggressive phenotypes, as reported in one NK-LGLL/CLPD-NK patient with a *STAT5B*^*N642H*^ mutation^11,59^. Besides the establishment of NK-cell leukemia, one N642H^NK/NK^ mouse developed a γδ T-cell leukemia. This is consistent with NKp46 expression on a subset of γδ T cells^60^ and the oncogenic potential of *STAT5B*^*N642*H^ in γδ T cells^27^. We observed a large population of GFP^+^ cells without CD3, TCRs, or NK1.1 on the surface in two diseased N642H^NK/NK^ mice. Upon transplantation, a GFP^+^ population expressing NK1.1 but not CD3 or TCRs accumulated in BM, spleen, blood and liver. We assume that these two aged mice developed an NK-cell leukemia but downregulated NK-cell lineage markers such as NKp46 and NK1.1, indicating a de-differentiated phenotype. De-differentiation has been observed in several tumor types^61^.

In addition to the findings from the N642H^NK/NK^ mouse model, the hypothesis that *STAT5B*^N642H^ promotes transformation of NK cells is further supported by the acquisition of cytokine independence in *STAT5B*^N642H^-transduced human NK-cell lines. This implies that *STAT5B*^N642H^ can drive uncontrolled growth and survival of NK cells independent of external cytokine signals, further contributing to the development and progression of NK-cell malignancies.

Indolent NK-cell leukemia/CLPD-NK may remain asymptomatic for an extended period – this aligns with the N642H^NK/NK^ model where most of the mice remain disease-free. In certain cases, it transforms into a more aggressive disease but the factors driving this transition are not understood. Our mouse model offers a tool to identify these factors, potential biomarkers and strategies to prevent the transformation.

Conventional therapeutic strategies against aberrant STAT5B signaling involve JAK inhibitors. There are currently more than ten different FDA approved JAK inhibitors available, and more are under investigation in advanced clinical trials^62^. Inhibition of JAKs has drawbacks as it affects additional signaling cascades leading to unintended side-effects. A more specific approach is the direct targeting of STAT5B using specialized STAT inhibitors or proteolysis targeting chimeras (PROTACs). The lack of an enzymatic activity in STAT5B and its close structural similarity to other STAT proteins complicates the development of specific compounds. While attempts are being made to develop such therapeutics, they have not yet advanced to the clinical stage^8,18,63^. This underscores the importance of identifying feasible therapeutic targets downstream of mutant STAT5B.

We identified 239 common *STAT5B* GOF DEGs between mouse and patient samples, with 136 of them significantly upregulated in STAT5B-mutated neoplasms. Pathway analysis revealed the TNFα/NF-κB signaling pathway significantly upregulated in mutant STAT5B samples. Aberrant levels of various cytokines and their receptors are often found in LGLL and contribute to the pathogenesis^64,65^. NF-κB activity is frequently upregulated in LGLL^64–66^. Likewise, *IL-7R* was identified as one of the top commonly upregulated genes. The interaction of IL-7R with its ligand, IL-7, can influence NK-cell maturation and function^67^. High IL-7R levels have been implicated in the tumorigenesis of T-cell leukemias and blocking the IL-7R inhibits the growth and survival of cancer cells^68–70^. We speculate that targeting TNF/NF-κB signaling or blocking the IL-7R would represent alternative therapeutic possibilities for NK-cell leukemia patients with *STAT5B* GOF mutations.

We recently identified a STAT5B regulated tetraspanin family member as a potential prognostic marker and therapeutic target in stem cell driven leukemias^21^. Among the top upregulated DEGs between mutant STAT5B murine and human samples, the tetraspanin *TSPAN13* stands out as a direct target gene of STAT5B^24,71^. TSPAN13 has been reported to facilitate cancer cell invasion and metastasis^72,73^. As the *TSPAN13* protein is exclusively detected in specific immune cell types such as plasmacytoid dendritic cells, B cells and NK cells (Human Protein atlas^74^), it represents an attractive target for NK-cell leukemia therapy. As a transmembrane protein, TSPAN13 represents a candidate for monoclonal antibody-based therapy to induce antibody dependent cellular cytotoxicity or as a target for CAR-NK/CAR-T-cell therapies. This targeted approach might minimize potential negative impact on other cell populations. Further evaluation is necessary to understand the significance of TSPAN13 in (STAT5B-driven) NK-cell leukemia development and therapy.

The conditional STAT5B^N642H^ mouse model provides a tool to investigate the effects of mutant STAT5B on specific cell types and its oncogenic/pro-tumorigenic potential. The transgenic STAT5B^N642H^ contains a V5-tag that facilitates the analysis of interaction partners and specific target genes through (chromatin) immunoprecipitation-based approaches. Our new mouse model represents a resource to investigate novel therapeutic interventions, as patients with *STAT5B*^N642H^ mutations often face therapeutic complications such as drug resistance or relapse^26,37^. Considering the urgent need for preclinical NK-cell neoplasm models, N642H^NK/NK^ mice will help us to understand NK-cell transformation and to identify novel biomarkers and therapeutic targets.

## Acknowledgements

We thank Sabine Fajmann, Petra Kudweis and Philipp Jodl for experimental support. We thank Bettina Wagner, Lill Anderson and Tina Bernthaler for their help in the generation of the new NK-cell specific STAT5B^N642H^ transgenic mouse model.

We thank the Next Generation Sequencing Facility at Vienna BioCenter Core Facilities (VBCF), member of the Vienna BioCenter (VBC), Austria. This work was supported by the Austrian Science Fund (FWF) Special Research Program SFB-F6107, the PhD program “Inflammation and Immunity” FWF W1212, Austrian Academy of Sciences doc.funds DOC 32-B28 and the FWF ZK-81B. The work was also supported by the Fellinger Cancer Research association and the Stadt Wien Kultur MA7 Grant. We thank Stephan Hutter (MLL) for bioinformatics support in the analysis of primary patient samples. BioRender.com was used for the graphical illustration of some Figures.

## Author Contributions

K.K., V.S. and D.G. conceived the study; T. R. and K.K. generated the mouse model. K.K., S.K., A.H., M.R, J.L., J.T., J.K. and D.G. performed the experiments. K.K., S.K. and D.G. analyzed the data. A.W.S. and B.M. established methods and helped with the experiments and analysis of the data. R.M. was involved in experimental design and scientific discussions; R.G., J.K. and S.K analyzed sequencing data; G.H. and W.W. provided bioinformatic patient data analysis; S.K., K.K., V.S., D.G. wrote the manuscript. D.G. and V.S. provided reagents and supervised the study; and all authors revised the manuscript.

