## Supplementary Material and Methods, Figures and Legends for "A lineage-specific *STAT5B*^N642H^ mouse model to study NK-cell leukemia"

This file includes the Supplementary Methods, Supplementary Figures S1-S6, Supplementary Figure Legends, the Supplementary References and an Annex Part I (uncropped western blot membranes).

#### Supplementary Materials and Methods

*Mouse crossing*

Homozygous R26-LSL STAT5B$^{N642H}$ knock-in mice (B6-*Gt(ROSA)26Sor*$^{tm1(STAT5B\text{-}N642H)}$) were crossed to *Vav*-Cre mice[1]. Their offspring carrying a heterozygous transgene integration at the R26 locus were used for the experiments.

All three homozygous R26-LSL knock-in lines (B6-*Gt(ROSA)26Sor*$^{tm1(STAT5B\text{-}N642H)}$, B6-*Gt(ROSA)26Sor*$^{tm2(STAT5B)}$ and B6-*Gt(ROSA)26Sor*$^{tm3(EGFP)}$) were crossed to *Ncr1*-iCreTg mice[2]. Mice carrying a homozygous integration of the transgenes have been used for all experiments with the R26-LSL lines crossed to *Ncr1*-iCreTg mice. Due to heterozygosity of *Ncr1*-iCre expression, the three mouse lines each contained Cre positive littermates (denoted as GFP$^{NK/NK}$ mice for the empty vector control line, STAT5B$^{NK/NK}$ mice for the line with the STAT5B transgene and as N642H$^{NK/NK}$ mice for the line with the STAT5B$^{N642H}$ transgene) and Cre negative littermates lacking transgene expression (denoted as “Cre neg”, which represents either a pool of the Cre-negative littermates from all the three different lines (GFP$^{NK/NK}$, STAT5B$^{NK/NK}$ and N642H$^{NK/NK}$ mice) or only N642H$^{NK/NK}$ mice).

Mice were maintained at the rodent facility of the University of Veterinary Medicine Vienna and the specific pathogen-free (SPF) quality of the animals was frequently confirmed according to FELASA recommendations for health monitoring[3]. Animal experiments were discussed and approved by the Ethics and Animal Welfare Committee of the University of Veterinary Medicine Vienna and the national authority (Austrian Federal Ministry of Education, Science and Research) in accordance with good scientific practice guidelines and national legislation, under licenses BMBWF-68.205/0103-WF/V/3b/2015, BMBWF-68.205/0010-V/3b/2019, BMBWF-68.205/0174-V/3b/20182022-0.762.012, 2023-0.108.862, 2022-0.404.452.

Aged mice as well as transplanted recipient mice were monitored daily for the onset of disease signs and sacrificed whenever reaching the pre-defined humane endpoints.

*Genotyping PCR*

PCR reactions were prepared on ice as follows: 1x Buffer II (Roche), 1 µM primer mix (R26_wt_rev, R26_wt_fw, R26_EGFP_fw), 2 mM dNTPs (Fermentas), 0.1 µl AmpliTaq DNA Polymerase (5U/µl), ddH2O per sample was added to a final of 20 µl reaction. The reaction was run at 95 °C (30 sec), 65 °C (30 sec) and 72 °C (40 sec) for 35 cycles on an Eppendorf Mastercycler. Each PCR product was mixed with 5 µl 6x DNA loading dye. 25 µl were loaded on a 1.5% (w/v) agarose gel immersed in 50x Tris Acetate (TAE) buffer until appropriate separation occurred. DNA was visualized with ethidium bromide in TAE Gel (1:20,000) exposed to ultra violet (UV) light in a Gel Doc™ XR Imager (Biorad). As reference marker the Gene RulerTM 1000 bp DNA ladder (Thermo Scientific) was used.

*Cell lines*

LentiX293T cells (Addgene) were cultured with DMEM (Sigma) complete medium, containing 10% FCS (Bio&Sell), 100 U/mL penicillin, 100 mg/mL streptomycin (Sigma), and 50 µM β - mercaptoethanol (Sigma).

Parental and transduced KHYG-1[4] (purchased from DSMZ) and IMC-1[5] (kindly provided by Prof. Chen, University of New Mexico Comprehensive Cancer Center) human NK-cell lines were cultured with RPMI complete with 100 U/ml, 25 U/ml or without human IL-2 (Proleukin, Novartis).

*Lentiviral transduction of human NK-cell lines*

Human non-mutant (WT) *STAT5B* or $STAT5B^{N642H}$ transgenes were cloned into a pWPI lentiviral expression vector (Addgene plasmid # 12254; kindly provided by the lab of Prof. Hoermann, previously at the Department of Laboratory Medicine, Medical University of Vienna), allowing for transgene expression coupled to IRES-controlled GFP expression. An empty vector control for expression of IRES-controlled GFP alone was included. Lentiviruses were generated by transfection of 4 µg of pWPI vectors (carrying the transgenes or the empty vector as a control) together with 3 µg of packaging vectors (2 µg of psPAX2 (Addgene plasmid # 12260) and 1 µg of pMD2.G (Addgene plasmid # 12259); kindly provided by Prof. Grebien, University of Veterinary Medicine Vienna) using Turbofect (Thermo Fisher Scientific) into around 80% confluent LentiX cells. 24h after the transfection, the medium was exchanged to RPMI complete medium containing 30 % FCS. After another 24h, supernatants were filtered through a 0.45-µm filter and used for the first round of lentiviral transduction. $3*10^5$ KHYG-1 or IMC-1 cells were resuspended in 1.5 ml of each of the lentiviral supernatants, supplemented with 100 U/ml IL-2 and 4 µg/ml Polybrene (Sigma), and seeded into 12-well plates. Spinoculation was performed for 90 minutes at 1000xg. After centrifugation, cells were incubated at 37°C and 5% CO2. On the next day, the transduced KHYG-1 and IMC-1 cells were washed and a second round of lentiviral transduction was performed with the lentiviral supernatants collected from the same LentiX cells (around 65-70h after transfection). After an overnight incubation at 37 °C, the remaining virus supernatant was washed away, and transduced KHYG-1 and IMC-1 cells were continued to be cultured in RPMI complete supplemented with 100 U/ml IL-2. To test how overexpression of WT STAT5B and the $STAT5B^{N642H}$ mutant impact on the growth of the transduced human NK-cell lines at a reduced IL-2 concentration, the transduced cell lines were cultured in parallel with 25 U/ml or 100U/ml

IL-2, starting from four days after the second lentiviral infection (4 days post infection (*p.i.*)). At 19 days *p.i.*, transduced cell lines that were cultured at 25 U/ml IL-2 were completely deprived of IL-2. The viability of the transduced human NK-cell lines was analyzed at multiple timepoints after IL-2 withdrawal by flow cytometry, using SYTOX$^{TM}$ Blue Dead Cell stain (Thermo Fisher Scientific) as a viability dye. Additionally, their percentages of GFP positivity were determined as a surrogate for transgene expression.

*Transplantation experiments*

For transplantation experiments from diseased N642H$^{vav/+}$ and N642H$^{NK/NK}$ mice, $5\times10^6$ splenocytes from diseased mice or control mice were injected intravenously (*i.v.*) into NSG (NOD.Cg-Prkdc$^{scid}$ Il2rg$^{tm1}$Wjl/SzJ) recipient mice.

Four transduced human NK-cell lines were selected in addition to the parental cell lines for transplantation into NSG mice. KHYG-1 +STAT5B$^{N642H}$ and IMC-1 +STAT5B$^{N642H}$ represent cells overexpressing the STAT5B$^{N642H}$ mutant that were obtained upon culture without IL-2, showing > 94 % transgene expression (evaluated by GFP positivity). The control cells overexpressing non-mutant STAT5B (KHYG-1 +STAT5B and IMC-1 +STAT5B) were sorted to achieve a > 98,6% transgene expression (evaluated by GFP positivity). The sorted KHYG1 +STAT5B cells were cultured at 100 U/ml IL-2, while IMC-1 +STAT5B cells could be maintained at 25 U/ml IL-2. $3*10^6$ or $1*10^6$ of these transduced human NK cell lines were injected *i.v.* into NSG recipient mice.

All transplanted mice were monitored daily for onset of disease symptoms and sacrificed once they reached the humane endpoints.

*Hematocytometry and Flow Cytometry*

Blood was collected into EDTA tubes (Greiner Bio-One Mini-Collect K3EDTA Tubes (Thermo Fisher Scientific)) from the V. *facialis* or via cardiac puncture under terminal anesthesia. White blood cell (WBC), lymphocyte (LYM), platelet (PLT) counts and haematocrit (HCT) were measured using an animal blood counter (scil Vet ABC).

For flow cytometric analysis, single cell suspensions were prepared from spleens, livers (LIV), bone marrow (BM), lymph nodes and lungs. Isolation of hepatic leucocytes was performed by separation with 37.5% percoll (GE Healthcare, Chicago, Illinois, USA). Lysis of erythrocytes from single cell suspension of BM, spleen and liver was performed with Red blood cell Lysis Buffer (150 mM NH4Cl, 10 mM KHCO3, 0.1 mM Na2EDTA, pH 7.2-7.4) prior to incubation with fluorescently labelled antibodies. For blood analysis, erythrocyte lysis was performed after the antibody staining.

Purified anti-CD16/CD32 antibodies (clone 93; Thermo Fisher Scientific (eBioscience™)) were used for blocking of Fc receptors.

Fluorochrome-conjugated antibodies (clones) directed against following proteins were purchased from ThermoFisher Scientific (eBioscience™): CD4 (GK1.5), CD8a (53-6.7), CD11b (M1/70), CD19 (eBio1D3), CD34 (RAM34), CD45.2 (104), CD122 (5H4), CD135 (A2F10), c-kit (2B8), Gr-1 (RB6-8C5), NKp46 (29A1.4), Sac-1 (D7), Ter119 (TER-119); from Biolegend/Biozym: CD3 (17A2), CD3e (145-2C11), CD25 (PC61), CD44 (IM7), CD48 (HM48-1), CD90.2 (53-2.1), CD150 (TC15-12F12.2), NK1.1 (PK136), TCRβ (H57-597), TCRγδ (GL3); or from Miltenyi Biotec: CD27 (REA499), KLRG1 (REA1016).

Analysis of apoptosis of splenic NK cells was performed *ex vivo* by staining with AnnexinV and 7-AAD using Annexin V Apoptosis Detection Kits (Thermo Fisher Scientific) according to the manufacturer’s protocol.

For intracellular staining of perforin (clone S1600A, Biolegend) and pSTAT5 (clone 47, BD), splenocytes were fixed with 2% paraformaldehyde (Sigma) after staining for surface markers and permeabilized with ice-cold 90% methanol before intracellular stainings. For detection of intracellular granzyme B (clone NGZB, Thermo Fisher Scientifc (eBioscience™)), BD Cytofix/Cytoperm™ Fixation/Permeabilization Solution Kit (BD Bioscience) was used according to manufacturer's instructions.

Total cell numbers were assessed by flow cytometry using Count Bright Beads (Invitrogen).

Flow cytometry samples were acquired using a Cytoflex (Beckman Coulter) or Cytoflex S (Beckman Coulter) flow cytometer. Data were analyzed using CytExpert (Beckman Coulter) or FlowJo software.

*Histology*

Organs were fixed overnight in 4% phosphate buffered formaldehyde solution (Roti®Histofix, Carl Roth, Karlsruhe, Germany), dehydrated, embedded and cut (4 μm). Paraffin-embedded sections were stained with hematoxylin and eosin (H&E) according to standard histological procedures. Blood smears were stained using a Hemacolor® Rapid staining of blood smears kit (Sigma, St. Louis, Missouri, USA). Images were taken using an Olympus IX71 microscope and CellSens Dimension Software (Olympus, Tokyo, Japan).

*NK-cell culture*

Splenic NK cells were isolated using DX5-labeled MACS beads according to the manufacturer’s instructions (Miltenyi Biotec). NK cells were cultured for up to 7 days in RPMI complete medium (10% FCS (Bio & Sell), 100 U/mL penicillin (Sigma), 100 mg/mL streptomycin (Sigma), 50 mM2-mercaptoethanol (Sigma)) supplemented with 400ng/ml murine (m)IL-2 (kindly provided by Peter Steinlein, Research Institute of Molecular Pathology (IMP)).

*Cytokine stimulation of splenocytes or cultured NK cells*

For *ex vivo* analysis of pSTAT5 levels by intracellular flow cytometry staining, splenocytes were stimulated with 10 ng/ml mIL-15 (PeproTech) for 20 min.

For analysis of pSTAT5 levels by immunoblotting, NK cells were either directly lysed after 7 days of IL-2 culture or lysed after 3h of IL-2 withdrawal with or without restimulation with 400 ng/ml mIL-2 and 50 ng/ml mIL-15 (PeproTech) for 20min.

*Immunoblotting*

Protein lysates from BM cells or IL-2 cultured NK cells were prepared in 1x Laemmli buffer, heated at 95°C for 5 min and sonicated at room temperature for 15 min. Protein concentrations were assessed using the Pierce™ BCA Protein Assay Kit (Thermo Fisher Scientific). Proteins were separated on 8-10% SDS polyacrylamide gels and transferred onto nitrocellulose membranes (Whatman®Protran®), which were blocked in 5% milk in pY-TBST buffer (10 mM Tris/HCl pH 7.4, 75 mM NaCl, 1 mM EDTA, 0.1% Tween-20). Membranes were incubated with antibodies against tyrosine-phosphorylated (pY-)STAT5 (Y694/699) (clone 47, BD Biosciences), total (t-)STAT5 (clone C-17, Santa Cruz or polyclonal IgG, R&D (cat. no. AF2168)), V5 (clone V5-10, Sigma Aldrich) overnight. β-Actin (clone AC-15, Santa Cruz) or α -Tubulin (clone 11H10, Cell Signaling Technology (CST)) served as loading controls. For detection of bound primary antibodies, membranes were incubated with horseradish peroxidase-conjugated anti-rabbit or anti-mouse antibodies (CST) followed by chemiluminescent imaging using Clarity Western ECL substrate (BioRad) and the ChemiDoc™ Touch Imaging System (BioRad). Image Lab software (BioRad) was used to process images and band intensity was quantified using the ImageJ Software.

### Supplementary Figures

#### Figure S1: N642H$^{vav/+}$ mice develop a hematopoietic malignancy

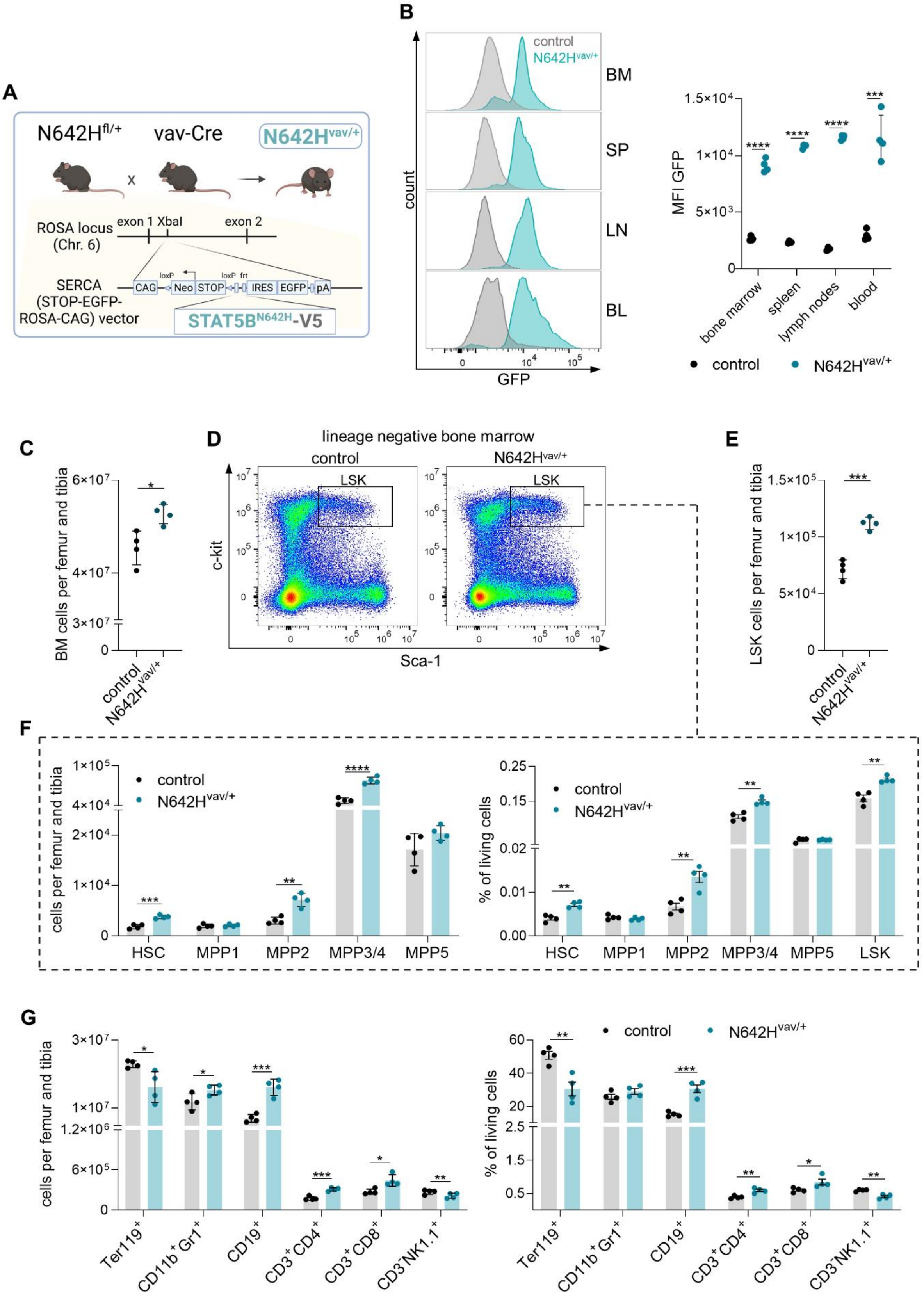

**Figure S1: N642H$^{vav/+}$ mice develop a hematopoietic malignancy**

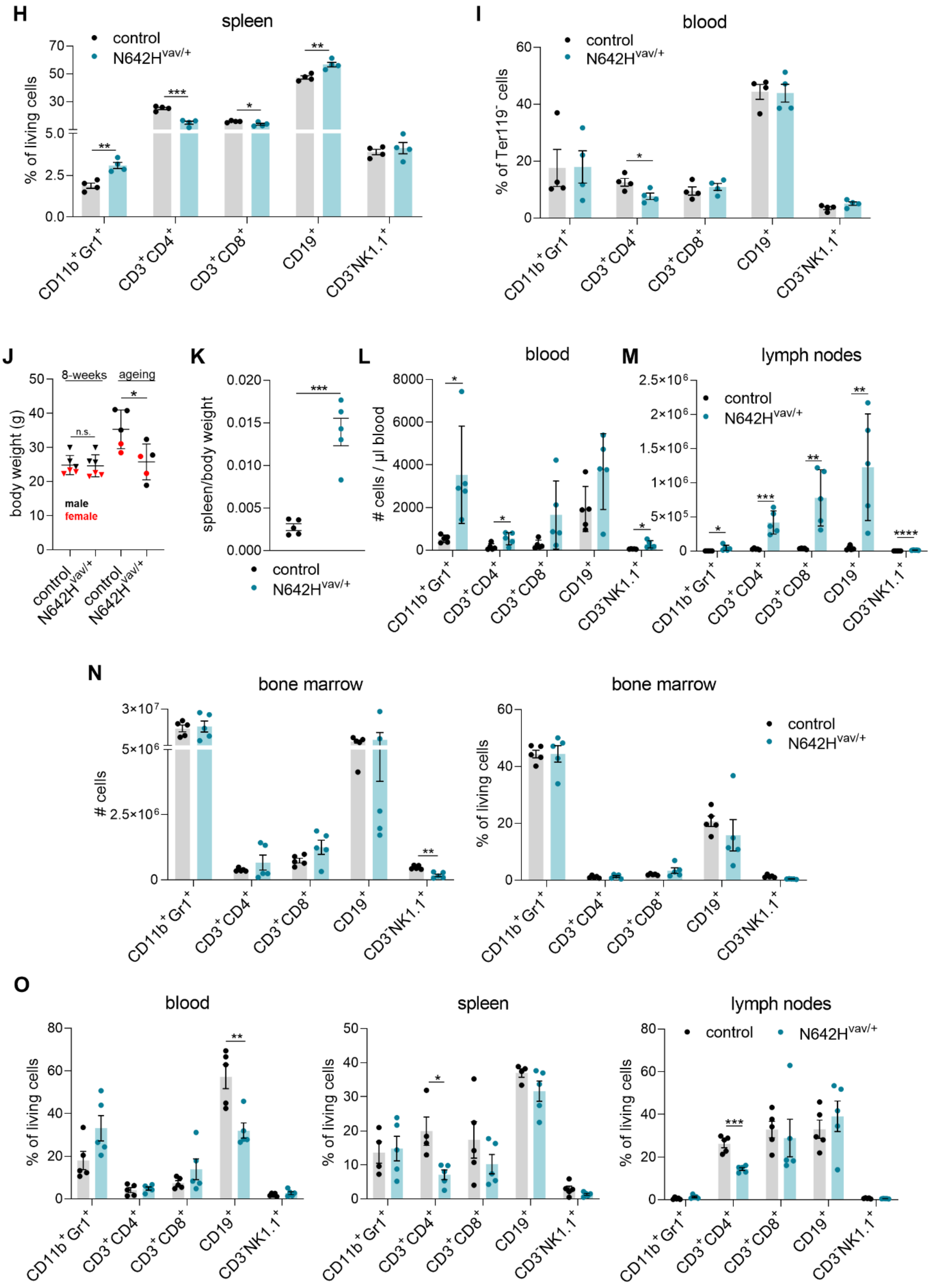

**Figure S1. N642H$^{vav/+}$ mice develop a hematopoietic malignancy**

(A) Schematic overview of the generation of N642H$^{vav/+}$ mice. (B) Flow cytometric analysis of GFP levels in different tissues of control and N642H$^{vav/+}$ mice. (*Left*) Representative histograms and (*Right*) quantification of GFP signals in BM, spleen, lymph nodes or blood derived from 8-week-old control and N642H$^{vav/+}$ mice (n=4/genotype, mean±SD). (C)-(G) Flow cytometry analyses of BM from 8-week-old control and N642H$^{vav/+}$ mice. Quantification of (C) total cell numbers (n=4/genotype, mean±SD), (D) representative gating of LSK population on lineage negative BM cells and (E) LSK cell numbers (n=4/genotype, mean±SD). (F) (*Left*) Total cell numbers and (*Right*) relative quantification of HSC and MPP1-5 cells (n=4/genotype, mean±SD). HSC subpopulations: HSC (LSK, CD34$^{-}$CD48$^{-}$CD150$^{+}$CD135$^{-}$), multipotent progenitor (MPP)1 (LSK, CD34$^{+}$CD48$^{-}$CD150$^{+}$CD135$^{-}$), MPP2 (LSK, CD34$^{+}$CD48$^{+}$CD150$^{+}$CD135$^{-}$), MPP3/4 (LSK, CD34$^{+}$CD48$^{+}$CD150$^{-}$). (G) (*Left*) Total cell numbers and (*Right*) relative quantification of Ter119$^{+}$ erythrocytes, CD11b$^{+}$Gr1$^{+}$ myeloid cells, CD19$^{+}$ B cells, CD3$^{+}$CD4$^{+}$-, CD3$^{+}$CD8$^{+}$ T cells and CD3$^{-}$NK1.1$^{+}$ NK cells (n=4/genotype, mean±SD). Relative quantification of CD11b$^{+}$Gr1$^{+}$ myeloid cells, CD3$^{+}$CD4$^{+}$-, CD3$^{+}$CD8$^{+}$ T cells, CD19$^{+}$ B cells and CD3$^{-}$NK1.1$^{+}$ NK cells in (H) spleen and (I) blood (n=4/genotype, mean±SD). (J) Body weight quantification of 8-week-old and aged control and N642H$^{vav/+}$ mice (n=5-6/genotype, mean±SD). (K) Relative quantification of spleen weights of aged control and N642H$^{vav/+}$ mice (n=5/genotype, mean±SD). Quantification of CD11b$^{+}$Gr1$^{+}$ myeloid cells, CD3$^{+}$CD4$^{+}$-, CD3$^{+}$CD8$^{+}$ T cells, CD19$^{+}$ B cells and CD3$^{-}$NK1.1$^{+}$ NK cells in (L) blood and (M) lymph nodes (n=5/genotype, mean±SD). (N) (*Left*) Total cell numbers and (*Right*) relative quantification of CD11b$^{+}$Gr1$^{+}$ myeloid cells, CD3$^{+}$CD4$^{+}$-, CD3$^{+}$CD8$^{+}$ T cells, CD19$^{+}$ B cells and CD3$^{-}$NK1.1$^{+}$ NK cells in BM (n=5/genotype, mean±SD). (O) Relative quantification of CD11b$^{+}$Gr1$^{+}$ myeloid cells, CD3$^{+}$CD4$^{+}$-, CD3$^{+}$CD8$^{+}$ T cells, CD19$^{+}$ B cells and CD3$^{-}$NK1.1$^{+}$ NK cells in (*Left*) blood, (*Middle*) spleen and (*Right*) lymph nodes (n=5/genotype, mean±SD).

Levels of significance were calculated using unpaired t-test in (B) – (O). *$p < 0.05$, **$p < 0.01$, ***$p < 0.001$ and ****$p < 0.0001$.

**Figure S2: *STAT5B*$^{N642H}$ drives the expansion of leukemic NKT-/ T cells upon transplantation**

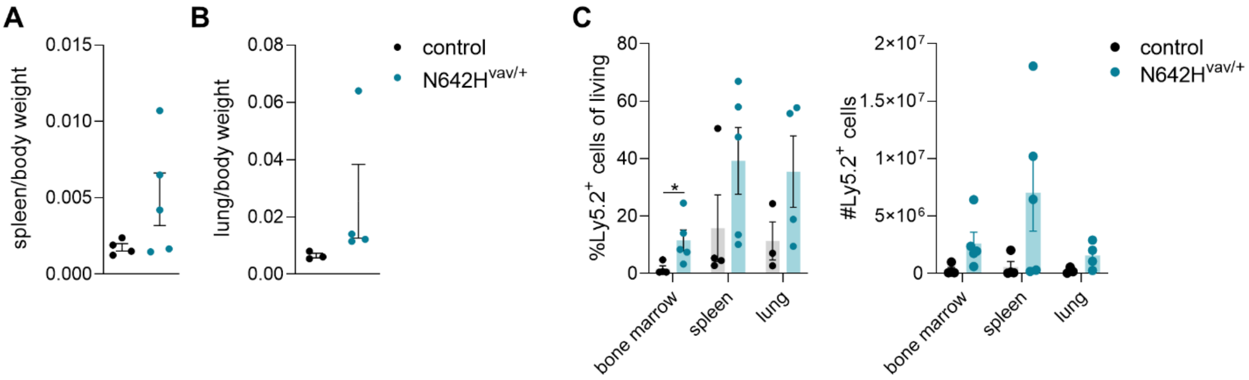


**Figure S2. *STAT5B*$^{N642H}$ drives the expansion of leukemic NKT-/ T cells upon transplantation**

Relative quantification of (A) spleen and (B) lung weights of NSG recipient mice injected with control or N642H$^{vav/+}$ cells (n=3-5/genotype, mean±SD). (C) (*Left*) Relative and (*Right*) total Ly5.2$^{+}$ cell numbers in BM, spleen and lung of NSG recipient mice (n=3-5/genotype, mean±SD).

Levels of significance were calculated using Mann-Whitney test in (C). *$p < 0.05$.

**Figure S3: *STAT5B*$^{N642H}$ promotes cytokine independence of leukemic human NK cells**

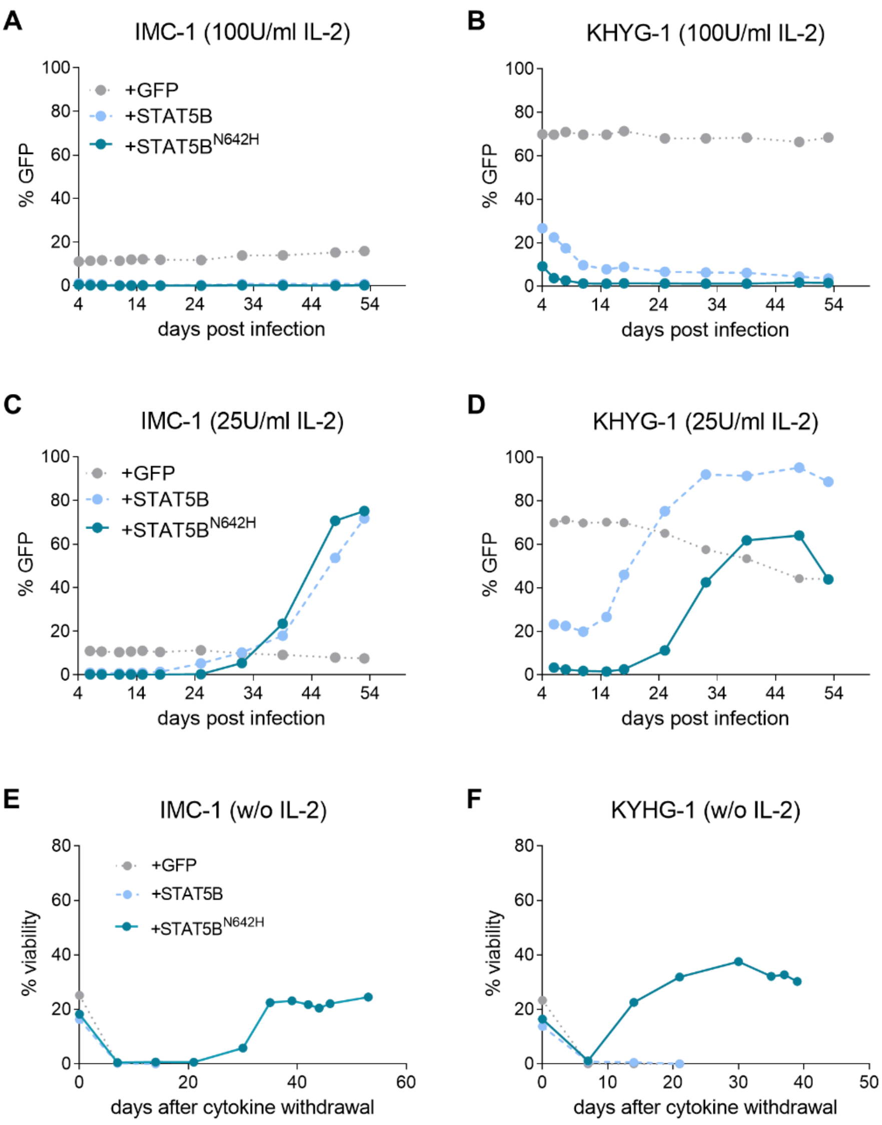


**Figure S3. *STAT5B*$^{N642H}$ promotes cytokine independence of leukemic human NK cells**
(A, C) KHYG-1 and (B, D) IMC-1 cell lines were transduced with empty vector (+GFP), non-mutant STAT5B (+STAT5B) or STAT5B$^{N642H}$ (+STAT5B$^{N642H}$). Four days after transduction, cells were either continued to be cultured with 100 U/ml IL-2 (A, B) or a reduced IL-2 concentration of 25 U/ml (C,D). (A-D) Percentage of GFP$^{+}$ cells was monitored over time. After complete IL-2 withdrawal, viability of transduced (E) KHYG-1 and (F) IMC-1 was analyzed.

**Figure S4: A NKp46⁺-cell specific mouse model to study the oncogenic role of *STAT5B*$^{N642H}$ in NK cells**

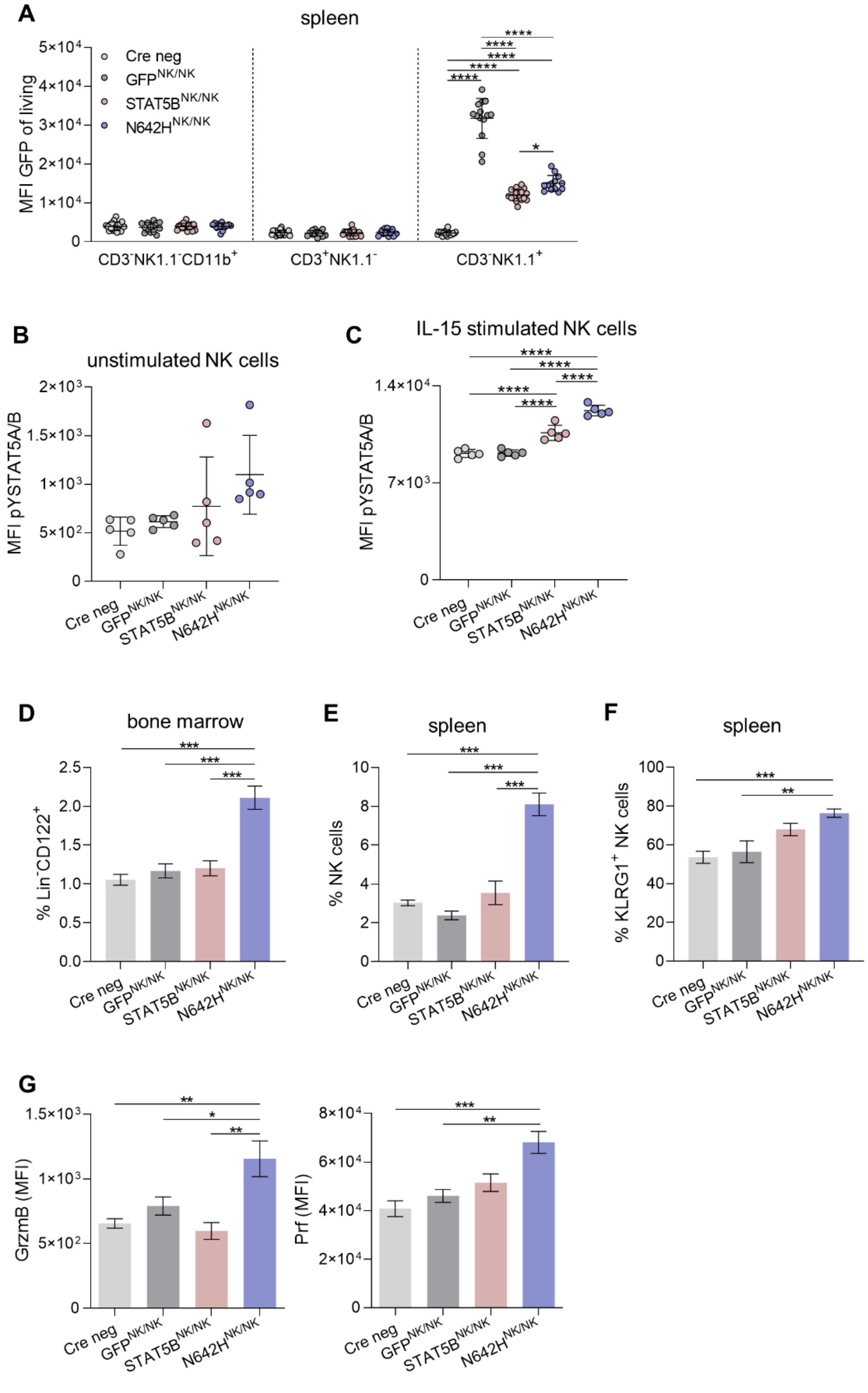


220

221

**Figure S4. A NKp46$^{+}$-cell specific mouse model to study the oncogenic role of *STAT5B*$^{\mathbf{N642H}}$ in NK cells**

(A) Quantification of GFP levels (MFI) in CD3$^{-}$NK1.1$^{-}$CD11b$^{+}$ myeloid cells, CD3$^{+}$NK1.1$^{-}$ T cells and CD3$^{-}$NK1.1$^{+}$ NK cells in the spleen of GFP$^{NK/NK}$, STAT5B$^{NK/NK}$, N642H$^{NK/NK}$ and Cre neg mice (n≥13/genotype, mean±SD). Flow cytometric analysis of *ex vivo* pY-STAT5 levels in (B) unstimulated and (C) IL-15 stimulated splenic NK cells (n=5/genotype, mean±SD). (D) Percentage of Lin$^{-}$CD122$^{+}$ cells in BM of GFP$^{NK/NK}$, STAT5B$^{NK/NK}$, N642H$^{NK/NK}$ and Cre neg mice (n≥10/genotype, mean±SD). (E) Percentage of splenic NK cells in GFP$^{NK/NK}$, STAT5B$^{NK/NK}$, N642H$^{NK/NK}$ and Cre neg mice (n≥10/genotype, mean±SD). (F) Percentages of KLRG1$^{+}$ NK cells in the spleens of GFP$^{NK/NK}$, STAT5B$^{NK/NK}$, N642H$^{NK/NK}$ and Cre neg mice (n≥10/genotype, mean±SD). (G) Steady-state levels of (*left*) granzyme B (GrzmB) and (*right*) perforin (Prf) in splenic NK cells from GFP$^{NK/NK}$, STAT5B$^{NK/NK}$, N642H$^{NK/NK}$ and Cre neg mice (n≥4/genotype, mean±SD).

Levels of significance were calculated using one-way ANOVA. *p < 0.05, **p < 0.01, ***p < 0.001, ****p < 0.0001.

#### Figure S5: *STAT5B*$^{N642H}$ induces NK-cell leukemia in mice

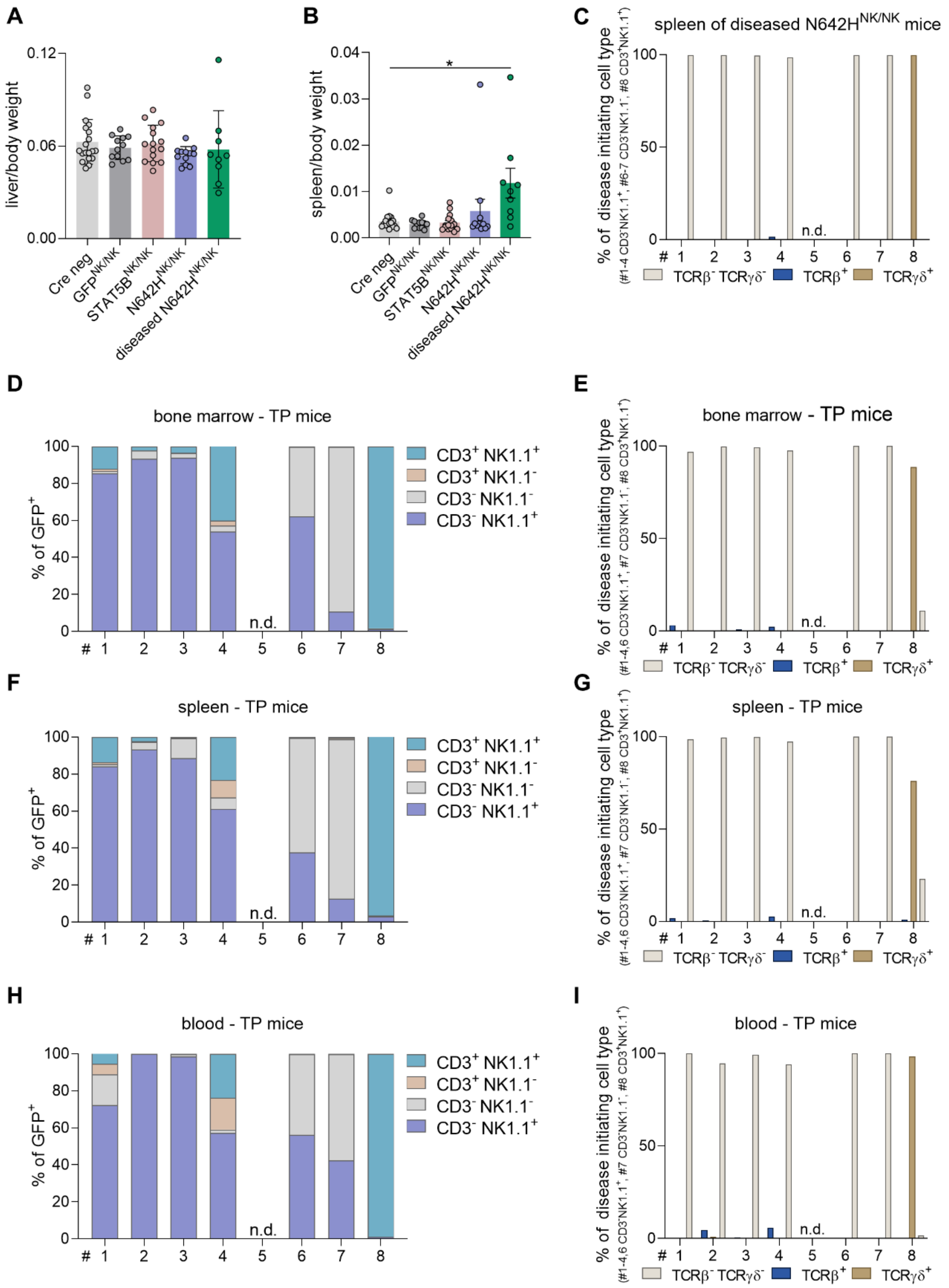


238

239

**Figure S5: *STAT5B*$^{N642H}$ induces NK-cell leukemia in mice**

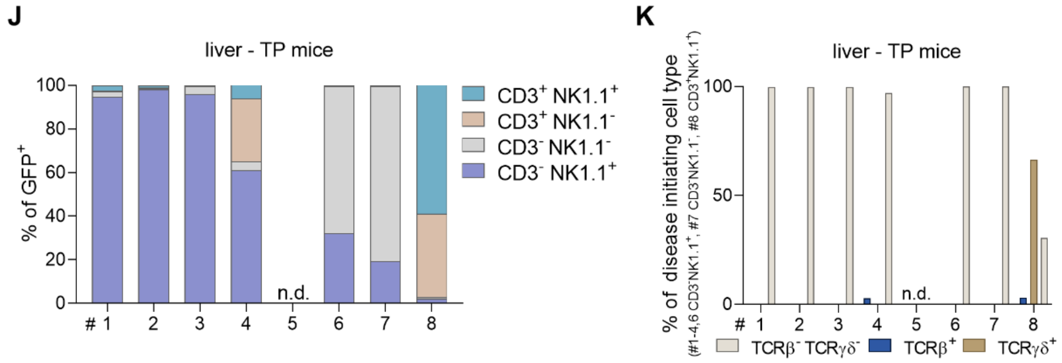


**Figure S5. *STAT5B*$^{N642H}$ induces NK-cell leukemia in mice**

Quantification of the (A) liver and (B) spleen to body weight ratio of Cre neg, GFP$^{NK/NK}$, STAT5B$^{NK/NK}$, N642H$^{NK/NK}$ and diseased N642H$^{NK/NK}$ mice (n≥8/genotype, mean±SD). (C) Relative quantification of TCRβ or TCRγδ expression on either CD3$^{-}$NK1.1$^{+}$, CD3$^{-}$NK1.1$^{-}$ or CD3$^{+}$NK1.1$^{+}$ splenocytes from diseased N642H$^{NK/NK}$ mice (n=7). Relative quantification of CD3$^{+}$NK1.1$^{-}$ T cells, CD3$^{+}$NK1.1$^{+}$ NKT cells, CD3$^{-}$NK1.1$^{+}$ NK and CD3$^{-}$NK1.1$^{-}$ cells in (D) BM, (F) spleen, (H) blood and (J) liver of diseased NSG recipients transplanted with diseased N642H$^{NK/NK}$ splenocytes (n=7). Relative quantification of TCRβ or TCRγδ expression on either CD3$^{-}$NK1.1$^{+}$, CD3$^{-}$NK1.1$^{-}$ or CD3$^{+}$NK1.1$^{+}$ cells in (E) BM, (G) spleen, (I) blood and (K) liver of diseased NSG recipients transplanted with diseased N642H$^{NK/NK}$ splenocytes (n=7).

Levels of significance were calculated using one-way ANOVA in (B). *p $<$ 0.05.

#### Figure S6: N642H$^{NK/NK}$ NK cells display molecular features of NK-cell leukemia patients harboring *STAT5B* GOF mutations

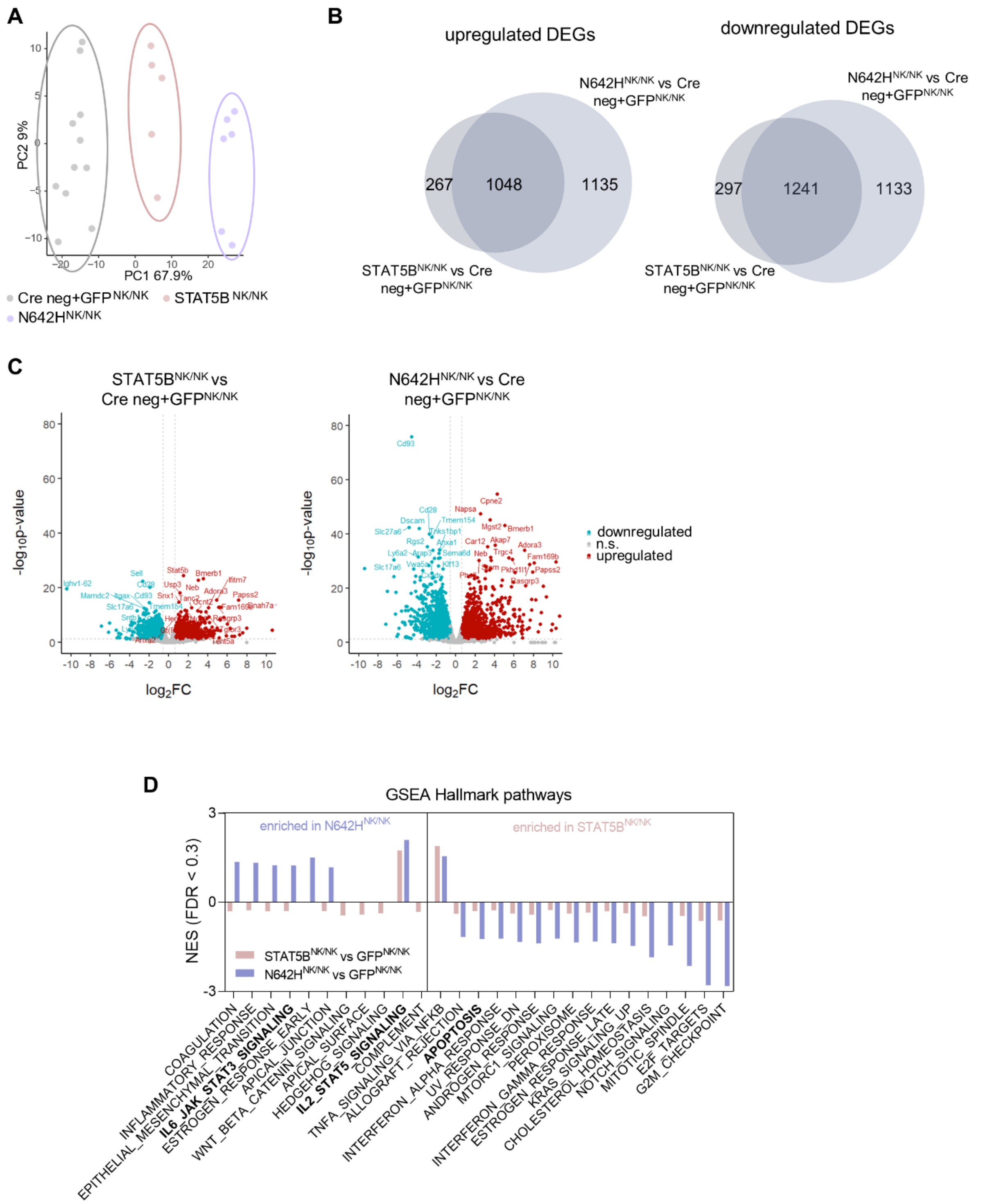

**Figure S6: N642H$^{NK/NK}$ NK cells display molecular features of NK-cell leukemia patients harboring *STAT5B* GOF mutations**

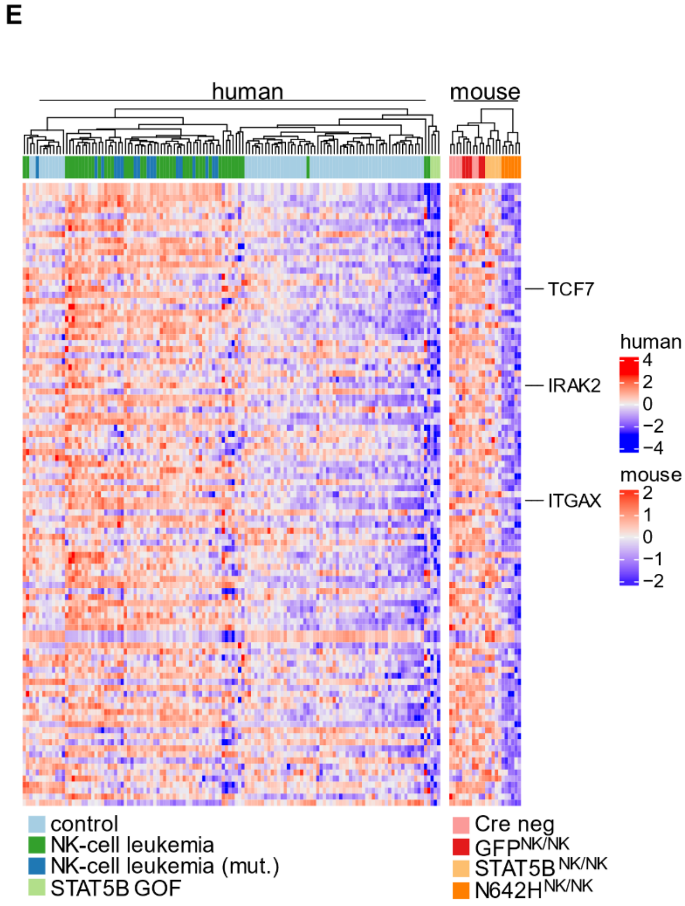


**Figure S6. N642H$^{NK/NK}$ NK cells display molecular features of NK-cell leukemia patients harboring *STAT5B* GOF mutations**

(A) PCA analysis of RNA-sequencing data of Cre neg (n=6), GFP$^{NK/NK}$ (n=5), STAT5B$^{NK/NK}$ (n=5), N642H$^{NK/NK}$ (n=6) NK cells. (B) Venn diagrams illustrating common up- or downregulated DEGs from STAT5B$^{NK/NK}$ (n=5) vs Cre neg (n=6) and GFP$^{NK/NK}$ (n=5) NK cells and N642H$^{NK/NK}$ vs. Cre neg (n=6) and GFP$^{NK/NK}$ (n=5) NK cells. (C) Volcano plots illustrating DEGs from STAT5B$^{NK/NK}$ (n=5) vs Cre neg (n=6) and GFP$^{NK/NK}$ (n=5) NK cells and N642H$^{NK/NK}$ vs. Cre neg (n=6) and GFP$^{NK/NK}$ (n=5) NK cells. (D) GSEA of HALLMARK pathways comparing STAT5B$^{NK/NK}$ (n=5) vs GFP$^{NK/NK}$ (n=5) and N642H$^{NK/NK}$ (n=6) vs GFP$^{NK/NK}$ (n=5) NK cells. (E) Heatmap illustrating the expression of the common downregulated genes (N642H$^{NK/NK}$ vs. STAT5B$^{NK/NK}$ and *STAT5B* GOF vs. NK-cell leukemia) on human control (n=64), NK-cell leukemia (n=44), NK-cell leukemia (mut., carrying a JAK/STAT mutation) (n=17), *STAT5B* GOF (n=3) samples and murine Cre neg (n=6), GFP$^{NK/NK}$ (n=5), STAT5B$^{NK/NK}$ (n=5), N642H$^{NK/NK}$ (n=6) NK cells.

#### Annex

#### Part I: Uncropped WB membranes

##### Figure 1A

3 8%-polyacrylamide gels have been loaded with the same lysates. The membranes has been cut at the 70 kDa protein ladder signal. The respective loading control for each blot is shown on the right.

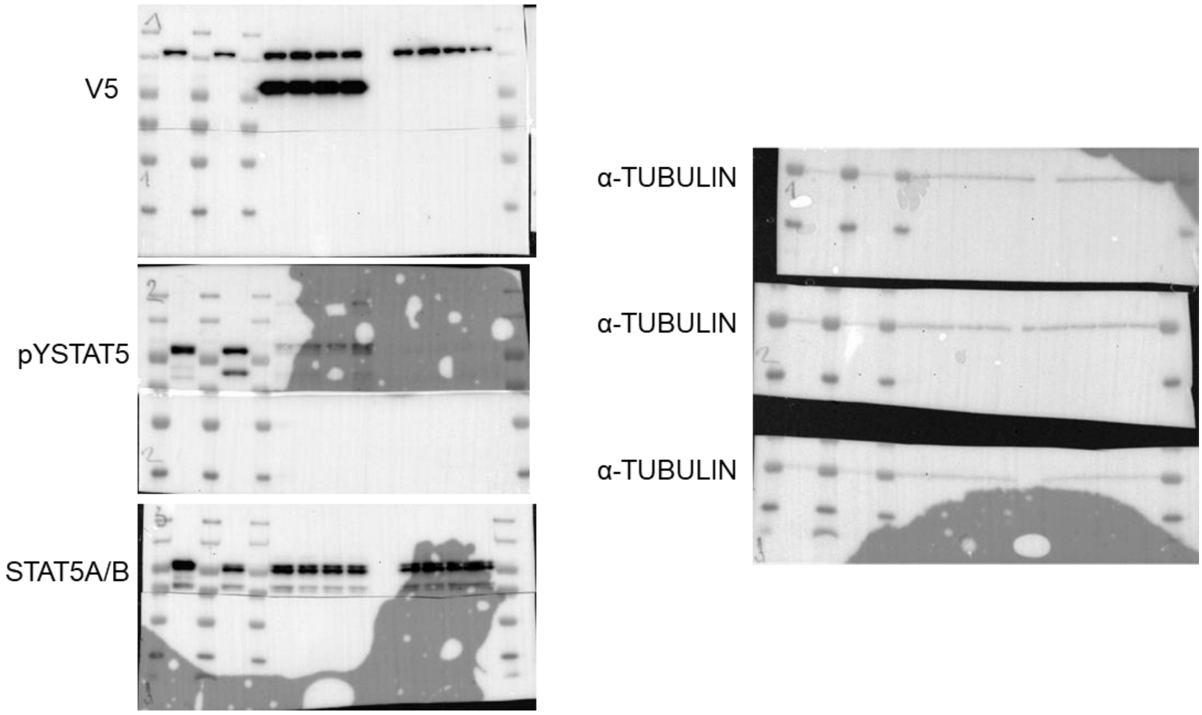
